# Transgressive and parental dominant gene expression and cytosine methylation during seed development in *Brassica napus* hybrids

**DOI:** 10.1101/2022.09.05.506610

**Authors:** Mauricio Orantes-Bonilla, Hao Wang, HueyTyng Lee, Agnieszka A. Golicz, Dandan Hu, Wenwen Li, Jun Zou, Rod J. Snowdon

## Abstract

The enhanced performance of hybrids though heterosis remains a key aspect in plant breeding, however the underlying mechanisms are still not fully elucidated. To investigate the potential role of transcriptomic and epigenomic patterns in early expression of hybrid vigour, we investigated gene expression, small RNA abundance and genome-wide methylation in hybrids from two distant *Brassica napus* ecotypes during seed and seedling developmental stages using next-generation sequencing technologies. A total of 71217, 773, 79518 and 31825 differentially expressed genes, microRNAs, small interfering RNAs and differentially methylated regions were identified, respectively. Approximately 70% of the differential expression and methylation patterns observed could be explained due to parental dominance levels. Via gene ontology enrichment and microRNA-target association analyses during seed development we found copies of reproductive, developmental and meiotic genes with transgressive and paternal dominance patterns. Interestingly, maternal dominance was more prominent in hypermethylated and downregulated features during seed formation. This contrasts to the general maternal gamete demethylation reported during gametogenesis in most plant species. Associations between methylation and gene expression allowed identification of putative epialleles with diverse pivotal biological functions during seed formation. Furthermore, most differentially methylated regions, differentially expressed siRNAs and transposable elements were found in regions flanking genes that had no differential expression. This suggests that differential expression and methylation of epigenomic features may help maintain expression of pivotal genes in a hybrid context. Differential expression and methylation patterns during seed formation in an F1 hybrid provide novel insight into genes and mechanisms with a potential role in early heterosis.

**Key message:** Transcriptomic and epigenomic profiling of gene expression and small RNAs during seed and seedling development reveals expression and methylation dominance levels with implications on early stage heterosis in oilseed rape.

## Introduction

Heterosis refers to the enhanced performance observed in F1 hybrids derived from two genetically distant, homozygous parents. The improved performance of hybrids compared to their inbred parents was described as early as the late 19^th^ century by Charles Darwin during his studies on maize and other plants (Darwin, 1876). Since Shull first coined the term “heterosis” in 1914 (Shull, 1914; 1948), this phenomenon has been applied in plant breeding to develop hybrids that outperform their inbred parents. However, despite the success of hybrid breeding in many major crops with selfing/outcrossing mating systems, for example maize, sunflower, tomato, sugarbeet and oilseed rape (Steeg et al., 2022), the mechanisms driving heterosis have yet to be fully elucidated. Several factors contributing to heterosis have been proposed: For extensive reviews see for example Wu et al. (2021) and Mackay et al. (2021). Classical quantitative genetics remains core to understanding heterosis as a product of allele interactions through dominance, over-dominance, or epistasis (Fujimoto et al., 2018). Nevertheless, through recent research, the cumulative understanding of molecular biology features has raised the question of how non-genetic and non-genomic features are also associated with heterotic patterns, and to which extent. Recent studies suggest that gene networks, allele bias, epigenomic and transcriptomic factors play a key role in heterosis (Wu et al., 2021; Yu et al., 2021).

*Brassica napus* (oilseed rape, canola; AACC, 2n=38) is not only an important crop where hybrid breeding has been implemented successfully, but is also a model crop of research interest due to its polyploid nature and phylogenetic proximity to *Arabidopsis thaliana*. Oilseed rape is the second most widely grown oilseed crop and has been the third most important food oil source worldwide in the last decade (FAO, 2022). The economic importance of hybrid breeding in oilseed rape is evident from the number and relevance of hybrid varieties in key producing countries like Canada, China and Germany. It is estimated that in 2015-2016 at least 80% of oilseed rape grown in China were hybrid varieties (Bonjean et al., 2016), whereas in Canada, the world’s largest producer of spring-type canola, herbicide-tolerant hybrid varieties have contributed to significant yield increases during the past decade (Malla and Brewin, 2019; FAO, 2022). The highest worldwide yields in winter-type oilseed rape in the past ten years were recorded in Germany (FAO, 2022), where the percentage of hybrid cultivars registered in the German National List increased from 74% in 2016 to more than 90% in 2022 (Friedt et al., 2018; BSA, 2022). These figures highlight the relevance of heterosis in current oilseed rape/canola breeding worldwide. The economic importance increases the need for a more refined understanding of the underlying molecular mechanisms behind heterosis in *B. napus*.

Technological advances in transcriptomic and epigenomic profiling in recent decades have increased awareness of regulatory and epigenetic factors in crop improvement and hybrid breeding (Scossa et al., 2021; Yang et al., 2021). “Omics” technologies not only help to describe and expand genetic diversity in crop species (Louwaars, 2018), but can also contribute to elucidating the role of regulatory and non-coding features in plants (Zanini et al., 2022). Transcriptomic and epigenomic features have been widely used to determine molecular and biological functions associated to improved performance in plant hybrids (Yu et al., 2021). For instance, RNA sequencing (RNA-Seq) data developed though microarrays and next-generation sequencing (NGS) has been used to find differential expressed genes (DEGs) linked to heterosis during diverse growth stages (Wang et al., 2017a; Zhu et al., 2020). Small RNAs (sRNAs) derived from endogenous genomic loci or exogeneous sources are known to regulate various functions and responses in plants. Classification and characterization of microRNAs (miRNAs) and small interfering RNAs (siRNAs) provide valuable information to investigate regulatory factors involved in modulation of gene and trait expression (Griffiths-Jones et al., 2006; Lunardon et al., 2020). For example, sRNAs have been associated with changes in performance in maize, rice and wheat (Zhang et al., 2014a; Li et al., 2014; Seifert et al., 2018a), while epigenomic features including chromatin interaction, histone modification and DNA methylation, can cause phenotypical changes without alterations in DNA sequences (Fitz-James and Cavalli, 2022). Genome-wide methylation differences in various plant species have been associated to phenotypic consequences (Muyle et al., 2022) and also linked to heterosis (Kawanabe et al., 2016; Lauss et al., 2018).

Differential gene expression studies in *B.napus* revealed key genes regulating flowering time, disease resistance and abiotic stress (Wu et al., 2016; Wang et al., 2017b; Jian et al., 2019), while small RNA profiling identified microRNA and siRNA sequences associated with pathogen response, abiotic stress and lipid metabolism in oilseed rape (Wang et al., 2017c; Jian et al., 2018; Martinez Palacios et al., 2019; Regmi et al., 2021). Furthermore, DNA methylation patterns were found to contribute to heat response, DNA repair and fertility in *B.napus* (Li et al., 2016; Ran et al., 2016; Wang et al., 2018; Yin et al., 2021).

Nevertheless, few studies have integrated multiple omics strategies to obtain a detailed scenario of expression and methylation patterns in oilseed rape (Shen et al., 2017; Wang et al., 2018). Interestingly, Shen et al. (2017) found specific expression and methylation patterns associated with heterosis in a commercial *B. napus* hybrid cultivar. Enhanced performance due to heterosis has been mostly evaluated at the genomic level and explained through allele interactions (Fujimoto et al., 2018) and introgressions of genomic regions between genetically and genomically distant parents (Hu et al., 2021a; Quezada-Martinez et al., 2021). Nevertheless, the transcriptomic and epigenomic networks involved in heterosis have not been fully elucidated, and the potential to include information on coding and non-coding features in hybrid breeding has been barely explored.

Comprehensive studies in maize and Arabidopsis demonstrated that heterosis can be observed in various developmental stages (van Hulten et al., 2018; Zhou et al., 2019). Heterosis during seed development can contribute directly to grain yield, seed biomass, germination and early vigour (Hochholdinger and Hoecker, 2007; Jahnke et al., 2010). Since seed formation is characterized by the merging of parental genomes, parent-specific epigenomic effects and genomic imprinting (Thiemann et al., 2009; Castillo-Bravo et al., 2022), it is an ideal stage for transcriptomic and epigenomic assessments in relation to heterosis. RNA-Seq and methylation-based studies have dissected putative heterotic loci in embryo and seed developmental stages in hybrids of Arabidopsis (Meyer et al., 2012; Kawanabe et al., 2016; Alonso-Peral et al., 2017; Chen et al., 2022). In *A. thaliana* and maize, early heterosis was associated to increases in cell size and number, seed yield and biomass (Jahnke et al., 2010; Wang et al., 2017a; Zhu et al., 2020). Groszmann et al. (2014) found that the maternal genotype was the major determinant of heterosis at early developmental stages in *A. thaliana*. Seed development is also well characterized for enriched epigenomic mechanisms through methylation and transcriptomic regulation, with pollen cells being hypermethylated and ovule cells demethylated in most plants (Batista and Köhler, 2020; Montgomery and Berger, 2021). Parental dominance effects are attributed with a key role during seed formation through diverging gamete methylation patterns (Weigel and Colot, 2012; Lauss et al., 2018). Moreover, the merging of parental genomes during embryogenesis can cause a genomic shock that can further alter the hybrid transcriptome (Bird et al., 2018).

Diverse studies in parent-offspring trios have compared parental dominant and transgressive gene expression patterns via expression level dominance (ELD) analyses in polyploids including *B. napus* (Yoo et al., 2013; Wu et al., 2018; Li et al., 2020) to elucidate the parental genotype effects on gene expression in interspecific hybrids between the diploid species progenitors. In the present study we analyse transcriptomic and epigenomic differences during seed and seedling development in homozygous parental lines and their F1 hybrid from a cross between the winter-type *B.napus* accession Express 617 (Lee et al., 2020) and the semi-winter accession *B. napus* G3D001 (Zou et al., 2018).

Preliminary observations showed significant heterosis in the hybrid between Express 617 × G3D001 in comparison to both parents under environmental conditions in central China. Numerous studies during the last decades has shown the heterotic advantages of crossing genetically distant oilseed rape varieties (Qian et al., 2009; Basunanda et al., 2010; Girke et al., 2012; Hu et al., 2021a). Hence, determining differentially expressed and methylated heterotic features between distant germplasm sources can potentially improve our understanding of the molecular mechanisms of heterosis. For this purpose, mRNA, sRNA and whole-genome bisulfite sequencing were carried in the two parental inbreds and their F1 hybrid. Differential features were identified and classified by their respective expression or methylation dominance levels to detect parental and hybrid-specific patterns associated with early developmental stages. Gene ontology enrichment (GO) and integration of omics features were performed to find putative interactions between the detected features and consequently evaluate their epigenomic and transcriptomic impact on early heterosis.

## Material and Methods

### Experimental design and growing conditions

Seeds from homozygous, advanced inbred lines of winter-type oilseed rape Express 617 (maternal line), semi-winter semi-synthetic oilseed rape G3D001 (paternal line) and their F1 hybrid offspring were planted in the 2020-2021 growing season at Huazhong Agricultural University of Wuhan field station. The third youngest leaf from each genotype were sampled from seedlings having six unfolded leaves (BBCH16) at 10 am under liquid nitrogen. Flower buds with similar sizes were selected on the fifth day after reaching full flowering (BBCH65) to perform manual selfing on a defined day in all genotypes along with crosses between Express 617 (female pollen recipient) and G3D001 (male pollen donor). The newly generated F1 crosses were employed to analyze the transcriptomic and epigenomic differences during seed formation between ovules pollinated from selfed F1 plants and those pollinated by outcrossing from Express 617 and G3D001 (referred to hereinafter as F0). Pollinated ovules were excised with forceps 15 and 30 days after pollination (DAP) at 10:00 am and immediately transferred to liquid nitrogen. Biological replicates consisted of pooled samples from the third youngest leaf from seven individual plants for leaf samples, and from four pollinated ovules from four different plants. The sampled tissue was aliquoted and used for all sequencing types described in this study. Moreover, three biological replicates were used for messenger RNA (mRNA) and small RNA expression experiments. Due to low material availability for some samples, only two biological replicates could be used for the methylation studies.

### mRNA, small RNA and whole genome bisulfite sequencing

mRNA was extracted using TRIzol™ Reagent (Thermo Fisher). A total of 0.5 μg of total RNA per biological replicate were used for preparing 150 bp paired-end (PE) read libraries using the NEBNext®Ultra™ II RNA Library Prep Kit (New England Biolabs, Inc.). Small RNA was extracted using a Plant miRNA kit (Omega Bio-tek Inc.). One microgram of total RNA per biological replicate was employed for the construction of 50 bp single-end (SE) reads using NEBNext^®^ Multiplex Small RNA Library Prep Set for Illumina™ (New England Biolabs, Inc.). Lastly, 2.5 μg of CTAB extracted-DNA per biological replicate were first treated with sodium-bisulfite using the Zymo EZ DNA Methylation-Lightning™ Kit (Zymo Research Corp.) and then built into 150 bp PE read libraries with the TruSeq Nano DNA LT Sample Prep Kit (Illumina Inc.) for whole-genome bisulfite sequencing (WGBS). All libraries were sequenced using an Illumina NovaSeq 6000 platform (Illumina Inc.). Read quality was evaluated with FastQC v.0.11.9 (Andrews., 2010) and multiqc v.1.9 (Ewels et al., 2016) for all sequencing types.

### mRNA and sRNA alignments

mRNA libraries were first filtered by selecting reads with an exact length of 150 bp, minimum base quality phred value of 5, no unqualified bases and less than 15% N bases using fastp v.0.23.1 -*q 5 -u 0 -n 15 -l 150* settings (Chen et al., 2018). Splice sites in the Express 617 reference genome assembly (Lee et al., 2020) were identified by first converting the gene annotation file format (Express617_v1_gene.gff3; MD5: cf26ec54823f348a0e23f027dc386a16) from a general feature format v.3 (GFF3) to a general transfer format (GTF) using the *agat_convert_sp_gff2gtf.pl* script from AGAT v.0.5.0 (Dainat, 2019). The output was then employed to find splice sites with the *hisat2_extract_splice_sites.py* script from HISAT2 (Kim et al., 2019). An index from the same Express 617 reference was built with *hisat2-build* function, and libraries were then aligned with HISAT2 using the *sensitive* preset and the *known-splicesite-infile* setting with the *hisat2_extract_splice_sites.py* previously generated file as input. Alignments were sorted and converted to a binary alignment map (BAM) format with samtools (Li et al., 2009) *view* and *sort* functions. The number of fragments in genes were counted with featureCounts 2.0.1 (Liao et al., 2014) using the AGAT GTF annotation file and the following settings *-p -B -C -Q 50 -t “exon” -g “gene_id”*, so that only read pairs having a minimum mapping quality of 50 and had both reads aligned to the same strand and chromosome were counted. Genes without any counts in all genotypes were removed. Small RNA libraries were first filtered by removing reads shorter than 18 bp with seqtk v.1.3 (Li, 2016). Subsequently, sRNA libraries were aligned against the Express 617 reference using ShortStack v.3.8.5 (Johnson et al., 2016). Only sRNA in which at least 80% of the primary reads had a length between 20-24 nucleotides, with less than 5 unpaired bases in secondary structure, and which were contained in predicted hairpin structures (i.e. only small RNAs clusters with *Y, N15, N14* or *N13* flags.) were considered as miRNA candidates. Small RNA sequences in which in which 80% of the primary reads had an exact length of 24 nucleotides and without miRNAs selection flags were regarded as putative siRNA. miRNA and siRNA clusters without any coverage in all biological samples were discarded prior to differential expression analysis.

### Expression level dominance analysis

Differential mRNA, miRNA and siRNA expression patterns between the F1 hybrid and its parents were assessed by comparing tissues within genotype trios in five tissues/stages: leaves from parental and F1 plants at six-leaf stage (BBCH16); ovules 15 days after pollination from selfed parents and F1 (OS15-F1) or F0 (OS15-F0); and ovules 30 days after pollination from selfed parents and F1 (OS30-F1) or F0 (OS30-F0), respectively. Differentially expressed genes, differentially expressed miRNAs (DE-miRNAs) and differentially expressed siRNAs (DE-siRNAs) between genotypes for each stage were identified using DESEQ2 (Love et al., 2014) with a padj value threshold < 0.05. The DESEQ2 built-in *estimateSizeFactors* and *counts* functions were used to extract the normalized counts which were then used for expression level dominance analyses. Briefly, student’s t-test (p < 0.05) from normalized counts of DEGs and DE-miRNAs identified in DESEQ2 were run between all genotypes for each comparison stage and gene. Tukey tests (p < 0.05) were then carried to rank each genotype by expression level. Finally, the resulting patterns were divided based on Yoo et al. (2013) as additive, dominant or transgressive. Gene expression heatmaps were generated with idep93 (Ge et al., 2018) using correlation distances and average linkages, and differentially expressed genes or sRNA shared between all stages were detected using the *Venn Diagrams* tool (VIB-UGent, 2021). In addition, the percentages of upregulated and downregulated DEGs from all genes per sugbenome, genotype and stage were calculated to evaluate subgenomic expression bias.

### Gene ontology enrichment

Gene models in the Express 617 reference assembly were functionally annotated through synteny comparison against the Darmor v.4.1 genome (Chalhoub et al., 2014) with inparanoid v.4.2 (O’Brien et al., 2005) using bootstrap, a BLOSUM80 (BLOcks SUbstitution Matrix) and an initial cut-off score of 60. Inparalogs with a similarity score equal or greater than 70 were selected for each gene. Pairs with only one homolog and with the highest similarity score were kept. The homologs were used for GO enrichment of biological processes based on expression level dominance for each stage, as well in comparisons between the F1 and F0 genotypes, using ShinyGo v.0.76 (Ge et al., 2020) with a 0.05 false discovery rate (FDR) cutoff. Only biological functions with more than one gene per biological pathway and with at least two GO groups were selected.

### DE-miRNA target prediction and mRNA interaction

Differentially expressed miRNAs sequences were extracted and used to predict their corresponding targets in Express 617 gene models using psRNATarget (Dai et al., 2018) with the version 2 scoring schema (Axtell, 2013). Maximum unpaired energy (UPE) of 25 and a flank length between 13 to 17 nucleotides in up/downstream region were set as target accessibility cutoffs. All possible targets for DE-miRNAs were reported since each miRNA can have multiple mRNA targets due to isomiRs formation. The DE-miRNAs were classified into putative miRNA families by blasting their sequences with BLAST (Altschul et al., 1990) against the mature miRNAs from the Brassicaceae family available at the miRBase sequence database release version 22.1 (Griffiths-Jones et al., 2006). Only the top five matches with the highest alignment scores and lowest expect values for each DE-miRNA were retained. Stem-loop sequences from the Brassicaceae family were used as BLAST targets when no mature miRNAs matches were found. Alternatively, if no Brassicaceae matches were found, then mature miRNAs and stem-loop sequences from the Viridiplantae clade were employed. The expression patterns from miRNA targets that were DEGs were compared with their associated targeting DE-miRNA expression to evaluate possible interactions between miRNA and mRNA target. The DEGs target functions were estimated by comparing their coding sequences against the Araport v.11 *Arabidopsis thaliana* coding sequences model (Cheng et al., 2017) via BLAST. Only the hit with the lowest expect value and not greater than 1.0 × 10^-4^, lowest identity percentage equal or above 90% and without gaps were selected.

### Bisulfite sequencing alignment and methylation level dominance

Reads with a minimum base quality phred value of 5, unqualified base percent limit of 50 and less than 15% N bases were selected from WGBS libraries using fastp v.0.23.1 -*q 5 -u 50 -n 15* settings (Chen et al., 2018). TrimGalore (Krueger et al., 2021) was then employed for trimming 8 basepairs from both 5′ and 3′ ends for each library as recommended for TruSeq libraries in the Bismark documentation. The Express 617 reference genome was bisulfite converted and indexed with Bismark v.0.23 (Krueger and Andrews, 2011) *bismark_genome_preparation* tool. Filtered reads were aligned to the bisulfite converted genome using *bismark* under default settings. Duplicates were afterwards removed with *deduplicate_bismark* and methylated cytosines (mC) were extracted using *bismark_methylation_extractor* while ignoring the first 2 basepairs from both 5’ and 3’ ends for both reads of a pair. Every mC in a CpG, CHG or CHH methylation context was selected and converted to a browser extensible data (BED) format with *bismark2bedGraph* using the --*cutoff 3 --CX* –and --*scaffolds* settings to select all nucleotides in which the methylation state was reported at least thrice.

The coverage for each mC in every methylation context was calculated with the *coverage2cytosine* from the Bismark package. The mC coverage in assigned chromosomes was then used as input for DMRCaller v. 1.22.0 (Catoni et al., 2018) to detect differentially methylated regions (DMRs). Each genotype within a trio was compared to each other using the *computeDMRs* function in 1000 bp bins with the *bins* method and the following settings: score test, a 0.01 p value threshold, and minimum cytosine count, methylation proportion difference and gap between bins of 4, 0.4 and 0 accordingly. The DMR methylation levels (i.e. the number of reads supporting methylation) were extracted from DMR output files and student’s t-test (p < 0.05) was run between all genotypes for each stage and DMR. Tukey tests (p < 0.05) were then used to rank the methylation within DMRs between genotypes and classified them by methylation level dominance (MLD) following the same categorization employed for ELD by Yoo et al. (2013). Shared and unique DMR across all stages were found with the *Venn Diagrams* tools (VIB-UGent, 2021).

### Cytosine methylation statistics and identification of methylated features

The number of mC nucleotides and the cytosine methylation level per 1 kbp bin (i.e. numbers of reads supporting cytosine methylation per bin) in each methylation context, genotype and stage were determined based on Bismark’s *coverage2cytosine* generated files using bedtools *makewindows* and *intersect* functions (Quinlan and Hall, 2010). In addition, DMRs were intersected with exons, introns, repeats and 1 kbp upstream promoter regions from Express 617 using bedtools *intersect* function. GO enrichment was analysed for differentially expressed genes having DMRs for all stages and genotypes. If no enrichment was detected, then the most frequent biological functions found in Ensembl Biomart (Cunningham et al., 2022) *B. napus* reference (Chalhoub et al., 2014) were reported. Detected differentially expressed genes having an additive or dominant expression level dominance pattern were defined as putative genetic epialleles if their loci coincided with corresponding additive or dominant methylation patterns in DMRs.

Heatmaps comparing the gene methylation and expression in transgressive DEGs were made with Heatmapper (Babicki et al., 2016) using Euclidean distances and average linkages to analyze the interaction between expression and methylation. Moreover, repeats in the Express 617 assembly were assigned to repeat families using RepeatModeler (Smit and Hubley, 2008) and CpG islands were identified with *cpgplot* from the EMBOSS v.6.6.0 package (Rice et al., 2000). CpG islands were called if the GC% was equal or greater than 50%, length greater than 200 bp and a minimum 0.6 observed to expected CpG dinucleotides ratio as described by Gardiner-Garden and Frommer (1987). Additionally, plots showing DEGs and methylation levels for each chromosome and stage, centromere loci and repeat density were made using the *circlize* package (Gu et al., 2014). Repeat density for each 1 kbp bin within each chromosome was calculated using bedtools while predicted Express 617 centromere loci were added based on Orantes-Bonilla et al. (2022).

Lastly, DMRs were intersected with DE-siRNAs, CpG islands and transposable elements (TEs) in 5 kbp upstream and downstream gene and DEG flanking regions in assigned chromosomes using bedtools to evaluate putative interactions between differentially methylated features and gene expression during seed development. The threshold was selected based on previous work on transposable elements and genomic imprinting in *B.napus* by Rong et al. (2021) and the fact that the average distance between genes in assigned chromosomes of the Express 617 reference is approximately 7.5 kbp. Chi-square tests followed by an FDR post-hoc adjustment (p < 0.05) were performed to find significant associations between differentially methylated and non-methylated features and determine their respective distances to genes or DEGs across all stages.

### Segmental expression assessment

Clustering of DEGs across chromosomal segments observed on *circlize* generated plots were further investigated. To assess the presence of expression clusters, segments that had more than 20 DEGs over a 500 kbp window were considered as putative differentially expressed segments. The threshold was selected on the basis that the Express 617 genome assembly has an average of 200 genes per 500 kbp and hence 20 genes would correspond to 10% of genes in the segment. The ratio of upregulated to downregulated DEGs per genotype and stage in each segment was calculated and normalized to Z-scores. Only segments showing clear differential patterns between genotypes based on Z-score heatmap clustering were retained. Such segments could either be a result of parental expression bias or due to commonly observed genomic rearrangements in allopolyploid *B. napus*. To investigate both possibilities, short read genomic sequence data from a G3D001 biological replicate was used for calling copy number variation (CNV) and investigating putative linkages between structural rearrangements and expression patterns. For this purpose, genomic DNA from a G3D001 ovule biological replicate taken 30 days after pollination was extracted using a CTAB protocol (Doyle and Doyle, 1987). Paired-end libraries were built with KAPA HyperPlus Kit (KAPA Biosystems) and sequenced with an Illumina NovaSeq 6000 platform (Illumina Inc.). Read quality was evaluated with FastQC v.0.11.9 and libraries were afterwards aligned with minimap2 (Li, 2018) against the Express 617 genomic reference (Lee et al., 2020). Alignments with both forward and reverse reads properly mapped (flags 99,163,147 and 83) were selected with samtools *view* and used to calculate coverage across chromosomes using the *bamtobed* and *genomecov* functions from bedtools. The coverage was used as input in a previously-described deletion-duplication pipeline (Stein et al., 2017), modified by excluding outliers if the depth was above 100 and by defining deletions and duplications as 25 kbp length segments that are one standard deviation above or below the mean coverage. Deletions and duplications were recorded in tab-separated files and intersected with differentially expressed segments using the *intersect* function in bedtools.

## Results

### Maternal dominant expression and methylation increases during seed development

Expression and methylation patterns from Express 617, G3D001 and their hybrid were compared during seed and seedling developmental stages. The parents were crossed during the experiment to evaluate developmental differences between selfed-F1 plants and Express 617 × G3D001 pollinated ovules that would develop into F1 plants (referred to here as F0). Figure 1 provides an overview of the plant materials, sampling tissues/timepoints and sample nomenclature. Next-generation sequencing yielded abundant coverage for each biological replicate (Tables S1-S3). Approximately 6.8 Gbp of mRNA sequences per biological replicate were aligned against the Express 617 genome assembly using HISAT2 v. 2.2.1 splice site aware aligner (Kim et al., 2019), producing mean alignment rates of 98.2% (Table S1). In addition, an average of 31 million sRNA reads per biological replicate were used to find putative miRNA and siRNA sequences with ShortStack v.3.8.5 (Johnson et al., 2016). Overall, each sRNA cluster had an average coverage depth of 186 (Table S2). Moreover, whole-genome bisulfite treated reads having a 31× genome coverage per biological replicate were aligned and processed with Bismark v.0.23 (Krueger and Andrews, 2011), as reported in Table S3.

**Figure 1.**
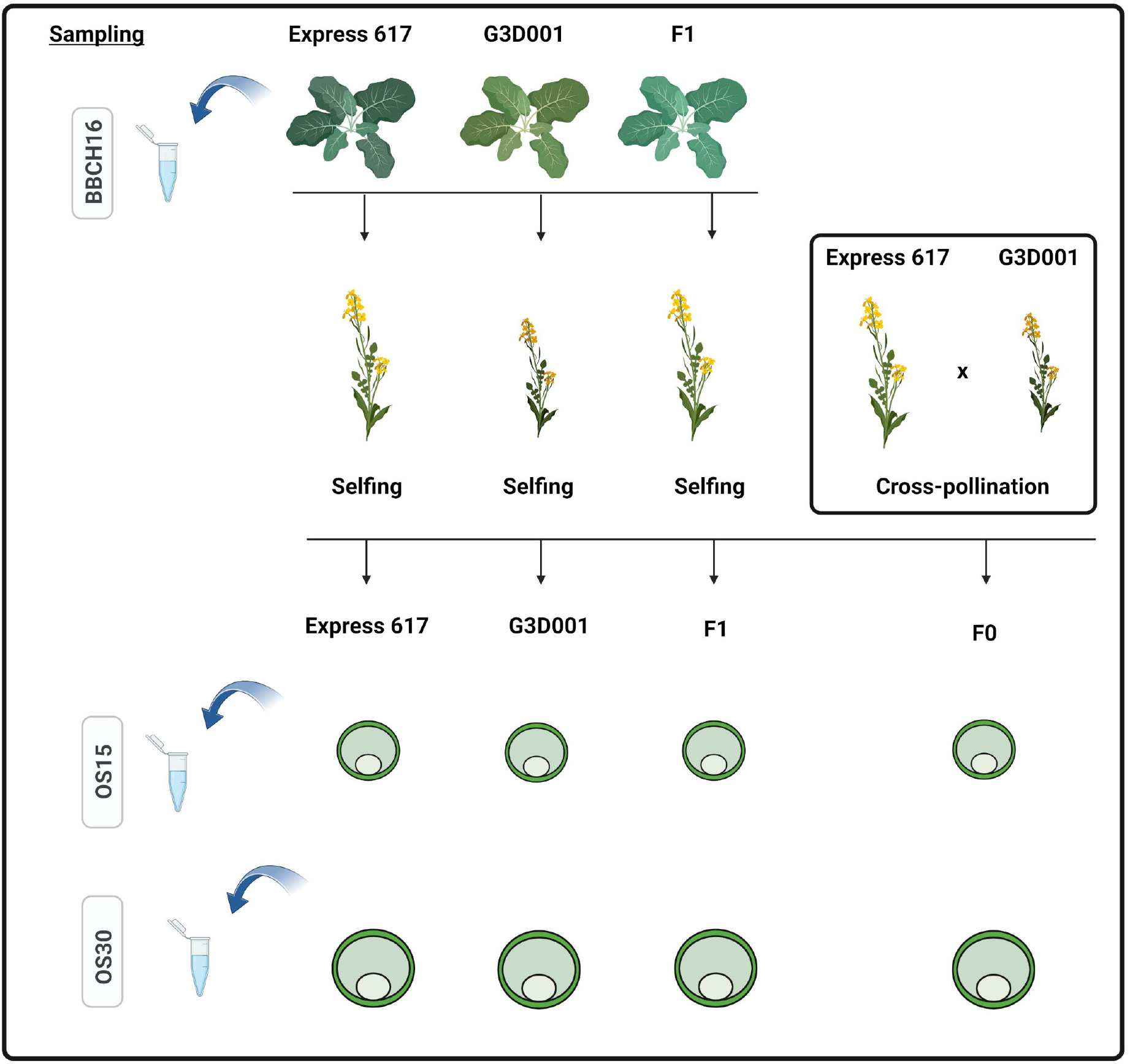
Transcriptomic and epigenomic experimental design. Leaves samples were taken at the six-leaves stages (BBCH16). Homozygous inbred plants of Express 617 and G3D001 along with their heterozygous F1 hybrid were self-pollinated to generate selfed ovules from each genotype. The two inbreed parents were also crossed during the experiment to develop cross-pollinated ovules (F0). Pollinated ovules were sampled and sequenced at 15 (OS15) and 30 (OS30) days after pollination.

All alignments were then employed to find features that were differentially expressed or differentially methylated between genotypes across all stages. In summary, a total of 71217 DEGs, 773 DE-miRNAs, 79518 DE-siRNAS and 31825 DMRs in both CpG and CHG methylation contexts were identified across all possible parents-hybrid comparisons per stage (Tables S4-S8). The detected features were evenly distributed across all chromosomes (Tables S9-S13). Differential features were further classified by their expression and methylation level dominance (Figure 2). More than 90% of the differentially expressed and methylated features corresponded to parental dominant and additivity models. Moreover, maternal dominance accounted for approximately 89%, 85%, 83% and 60% from all detected DEGs, DE-miRNAs, DE-siRNAS and DMRs in the F0, respectively, whereas paternal dominance was more prevalent in the F1-selfed offspring (Table S14). Furthermore, most maternal dominant DMRs in the F0 were hypermethylated, whereas DEGs were downregulated. This contrasts to the expected female gamete demethylation observed in seed formation in other plants (Batista and Köhler, 2020).

**Figure 2.**
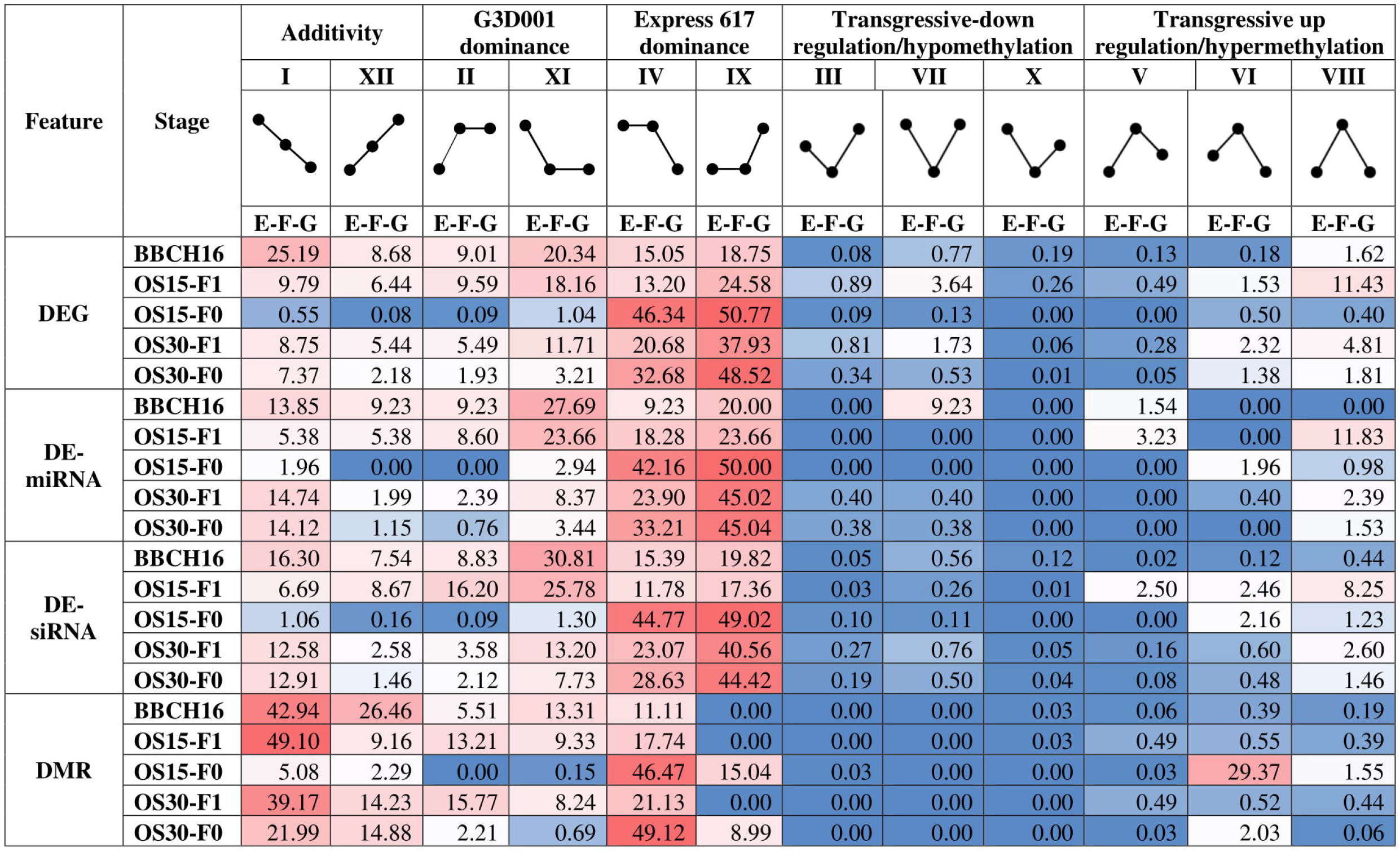
Percentages of differentially expressed genes (DEGs), differentially expressed miRNAs (DE-miRNAS), differentially expressed siRNAs (DE-siRNAS) and differentially methylated regions (DMRs) in CpG and CHG methylation contexts by expression level dominance (ELD) and methylation level dominance (MLD) patterns per stage. Increase and decrease in expression and methylation per patterrn are displayed by the dot-ended lines showing the relative expression or methylation levels for the parental genotypes Express 617 (E) and G3D001 (G) along with their F1 hybrid (F). Differential expression and methylation are displayed for leaf samples at stage BBCH16 and for ovules at 15 (OS15) and 30 (OS30) days after pollination by selfing (F1 ovules) or cross-pollination between the two parental lines (F0 ovules). Percentages are displayed with colored backgrounds to represent high (red) or low (blue) abundance.

Transgressive upregulated features, in which the hybrid has a higher expression than the parents, were more frequent in seeds from selfed-F1 plants compared to those from the recently formed F0. Maternal dominance from Express 617 accounts for most of the DEG and DE-siRNAs patterns observed, suggesting a potential maternal relevance in seed development. Interestingly, no gene expression bias was found between the A and C subgenomes (Table S15, Figure S1-S5); nevertheless, more upregulation was observed in the paternal line, while the maternal line displayed more downregulation during seed development. This contrasts with the expected gene silencing in the maternal genome that is attributed to maternal demethylation during seed formation. Moreover, a slightly higher number of differentially expressed features following maternal expression patterns were found in the F0 than in the selfed-F1; however, this might be due to the allele segregation in the selfed-F1 plants that would lead to the maternal parent being heterozygote and putatively reducing the number of features with maternal dominant expression.

A total of 1565 DEGs, 12 DE-miRNAs, 1111 DE-siRNAs, 896 DMRs in CpG context and 650 DMRs in CHG context were consistently detected across all stages (Tables S16). Altogether, differential features shared across all stages and genotypes corresponded to 3% of all detected features, whereas differential features unique to a single stage compromised approximately 2% of all detected features (Table S16). Differential features with consistent dominance level patterns across sampling stages are presented in Fig. 3. Interestingly, features with consistent dominance patterns in a higher number of stages tended to exhibit maternal dominance. These features were mostly shared between early and late pollinated ovule stages in the F1 and F0 (Table S17-S20), suggesting dominance of the maternal genotype during seed formation for both genotypes. Parental effects have been reported to play a role in heterosis in maize and Arabidopsis (Ma et al., 2018; Castillo-Bravo et al., 2022) and are further addressed in the discussion.

**Figure 3.**
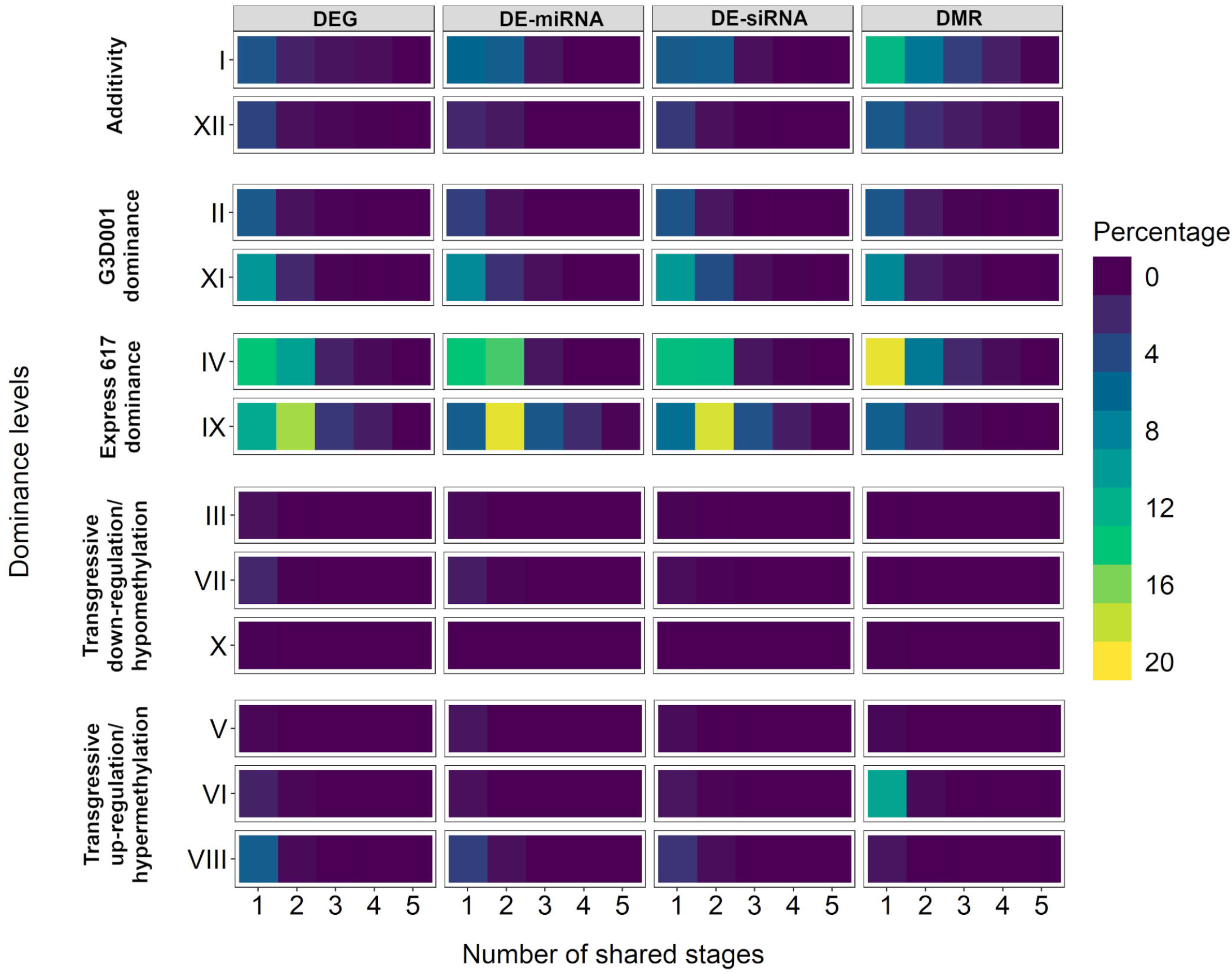
Percentage of shared differential features between stages based on dominance level patterns displaying differentially expressed genes (DEGs), differentially expressed miRNAs (DE-miRNAs), differentially expressed siRNAs (DE-siRNAs) and differentially methylated regions (DMRs) in CpG and CHG contexts.

### DEGs and differential miRNA expression regulate F1 seed development

Gene ontology enrichment for biological processes was carried out for all stages based on their expression level dominance. Significant enrichment for pivotal biological functions such as amino acid and carbohydrate synthesis, photosynthesis, protein transport and DNA repair and replication were found in 15 and 30 days after pollination ovules (Table S21). No significant enrichment was identified in leaves during the seedling stage. Only transgressively upregulated genes in F1 ovules after 15 days of pollination displayed terms associated with reproduction and meiosis (Figure 4, Table S22, Figures S6-S10). Differential gene expression and gene ontology between the F1 and F0 at 15 days after pollination showed an increase in photosynthesis-related functions in the F1 hybrid, whereas the F0 showed increased accumulation of energy reserve compounds and cell mobilization (Table S21). GO terms linked to carbohydrate metabolism, photosynthesis, stress response and cell development have been linked to heterosis in maize, rice, sunflower and oilseed rape (Bao et al., 2005; Lai et al., 2006; Ma et al., 2018; Zhu et al., 2020).

**Figure 4.**
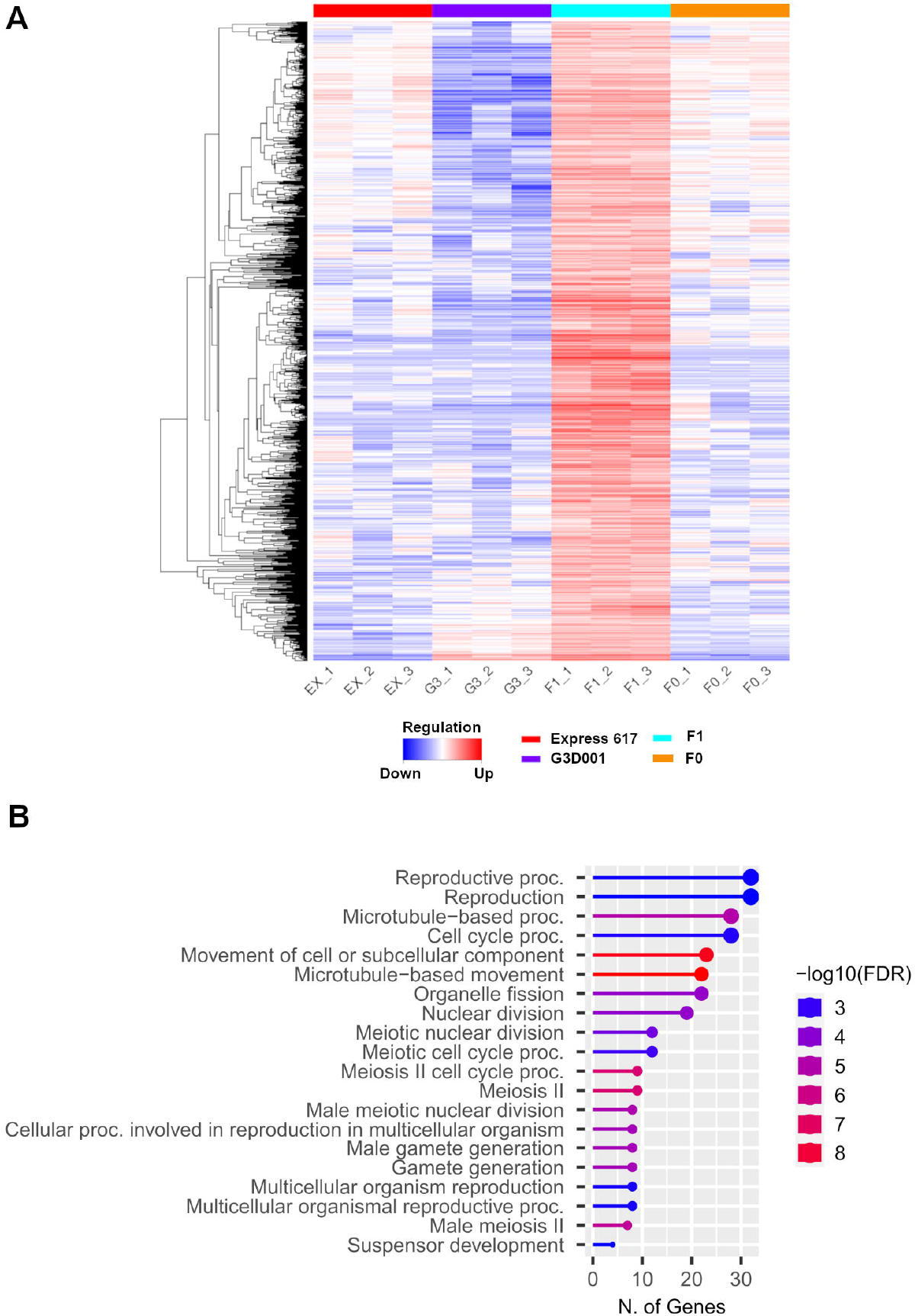
(**a**) Gene expression heatmap and (**b**) gene ontology (GO) enrichment of biological processes from 15 days after pollination ovules with transgressive upregulation patterns in the F1.

Additionally, 51 putative mRNA targets from all DE-miRNAs were detected across all stages (Table S23). Interactions between DE-miRNAs and DEG mRNA targets are reported in Table 1 and Table S24. Most DE-miRNAs associated with DEG targets had downregulated expression in the parents and F1 (ELD IX) and were more abundant during the late seed developmental stage. Expression from *B. napus* orthologs of *PHABULOSA (PHB)*, *REVOLUTA* (REV) and *TARGET OF EARLY ACTIVATION TAGGED 2* (TOE2), which are involved in plant growth and development, was not increased despite the low expression of miRNAs known to target these genes. Likewise, positive proportional expression interactions were observed in a *B. napus* ortholog of *EMBRYO DEFECTIVE 2204* (*EMB2204*) on chromosome A02, whereas an inversely proportional relationship between the miRNA and mRNA target was found for a *B. napus* ortholog of *EMBRYO DEFECTIVE 2016* (EMB2016)on chromosome A03 (Figure 5). Both genes are involved in embryo development, yet they appear to be regulated in an opposite manner, although further research is required to elucidate their role in *B. napus* seed formation. *PHB*, and possibly *PHAVOLUTA (PHV)*, are positive regulators of the *LEAFY COTYLEDON 2 (LEC2)*, a well-known regulator of seed maturation (Tang et al., 2012). A BLAST search from the *A. thaliana* Araport 11 assembly (Cheng et al., 2017) for the *LEC2* coding sequence (*AT1G28300.1*) revealed a single hit that passed the filtering criteria specified for miRNA targets in Materials and Methods. The ortholog corresponded to the *C05p022870.1_BnaEXP* gene model in the Express 617 genome assembly (Lee et al., 2020), which was found to be differentially expressed in late seed development (Table S17 and S24). This example further highlights the broader and indirect impact of miRNAs through gene network interactions. Such transcriptomic networks not only play critical roles in heterosis (Wu et al., 2021) but are also regulated partially by miRNAs (Dong et al., 2022).

**Table 1.** Predicted mRNA target from differentially expressed miRNAs (DE-miRNAs) in 15 and 30 days after pollination ovules in F1 and parents by expression level dominance (ELD).

| Stage | Predicted<br>DEmiRNA familiy | DEmiRNA-ELD | miRNA-Target | miRNA-target-ELD | *A. thaliana*<br>homolog ID | *A. thaliana*<br>homolog name |
| --- | --- | --- | --- | --- | --- | --- |
| **OS15-F1** | miRNA 165/166 A<br>miRNA 165/166 B<br>miRNA 165/166 C | IX | C04p041520.1_BnaEXP | IX | AT2G34710 | PHABULOSA |
| **OS30-F1** | miRNA 3629 A | VIII | A02p005120.1_BnaEXP | VIII | AT1G22090 | EMB2204 |
|  | miRNA 166 A<br>miRNA 165/166 B<br>miRNA 166 B<br>miRNA 166 C | IX | C04p041520.1_BnaEXP | IX | AT2G34710 | PHABULOSA |
|  | miRNA 166 A<br>miRNA 165/166 B<br>miRNA 166 B<br>miRNA 166 C | IX | C05p024820.1_BnaEXP | XII | AT1G30490 | PHAVOLUTA |
|  | miRNA 9410/9411 A<br>miRNA 9410 A | IX | A06p017700.1_BnaEXP | VII | AT3G47060 | FTSH PROTEASE 7 |
|  | miRNA 165/166 B | IX | A02p009610.1_BnaEXP | IX | AT5G60690 | REVOLUTA |
|  | miRNA 169 A | IX | A03p029900.3_BnaEXP | XI | AT3G05680 | EMB2016 |
|  | miRNA 172 A | IX | C09p033660.1_BnaEXP | IX | AT5G60120 | TOE2 |

**Figure 5.**
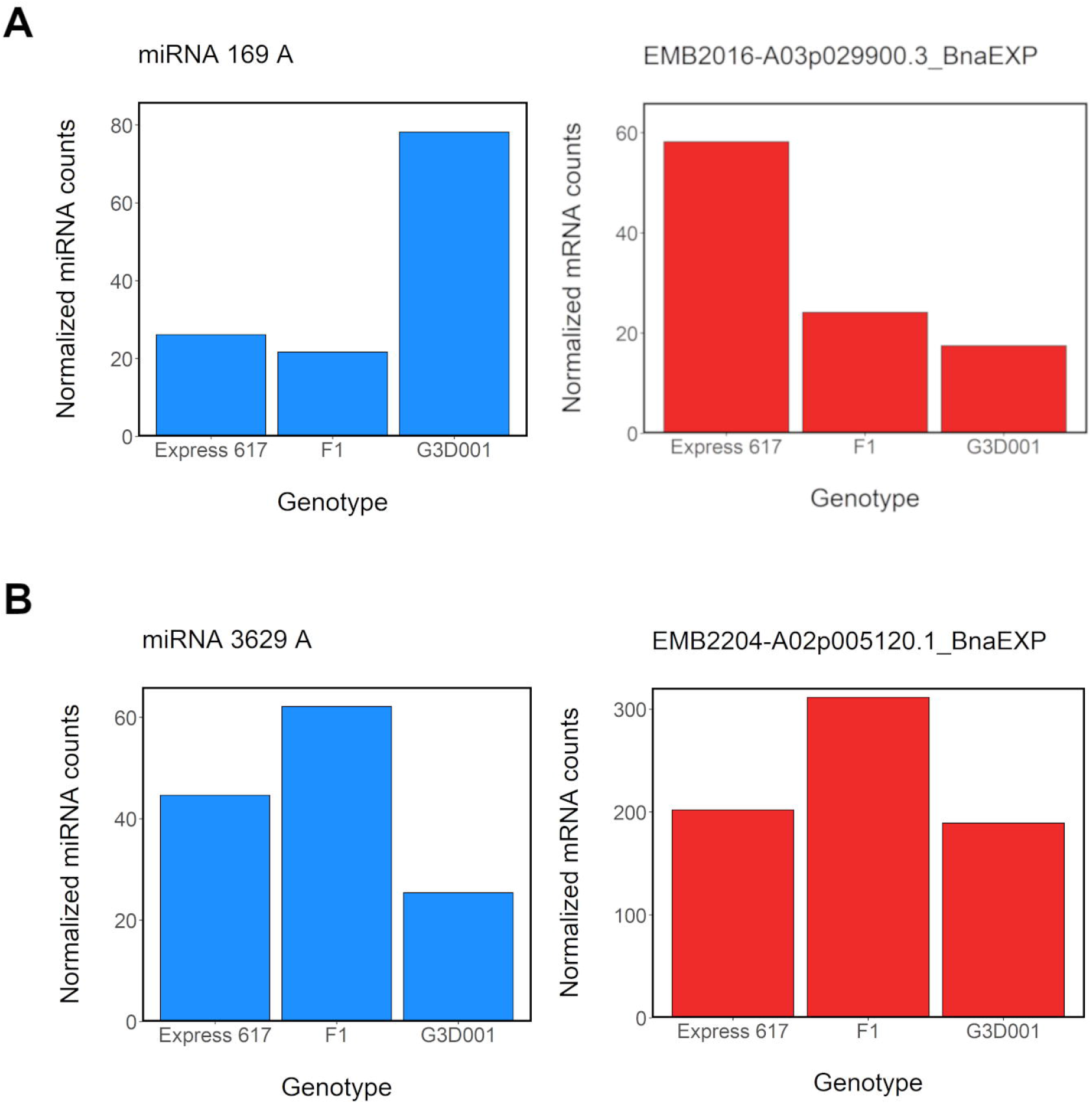
Normalized expression levels from selected differentially expressed miRNAs (DE-miRNA) and their respective differentially expressed target genes (DEG) in ovules 30 days after pollination in the F1 and parental genotypes, respectively. **a** Inversely proportional miRNA-mRNA target expression of miRNA 169A and a *B. napus* ortholog of its target gene *EMB2016* on chromosome A03 (A03p029900.3_BnaEXP). **b**: Proportional miRNA-mRNA target expression of miRNA 3629A and a *B. napus* ortholog of its target gene *EMB2204* on chromosome A03 (A02p005120.1_BnaEXP).

### Methylated features in early seed formation

Methylation levels were highest in the CpG context, with an average of 80% across all stages and genotypes (Fig. 6). Methylation levels in CHH context were the lowest, ranging from 20% to 27% despite having the highest number of methylated cytosines (Table S25, Fig S11-S14). No methylation bias per chromosome was observed (Table S25). Approximately 12%, 14% and 10% of DMRs were in promoters, exons and introns, respectively, whereas a high percentage of DMRs (43%) were inside repeat motifs (Table S26). Repetitive sequences account for 37.5% of the Express 617 genome (Lee et al. 2020), and although 66% of repeats were methylated with an average 41% methylation level, less than 1% were differentially methylated (Table S27). Most differentially methylated transposable element (TE) families and superfamilies coincided with those that are most frequent in the Express 617 reference genome, such as LTR (long terminal repeat) Copia and Gypsy families. Approximately 70% of these were found within 5 kbp flanking regions of genes (Table S28-S29). Chi-square tests followed by FDR adjusted post-hoc testing at p > 0.05 revealed a significant association between the analyzed genomic features (DMRs, TEs, and differentially methylated or non-methylated DE-siRNAs) in terms of their distance to genes and DEGs (Table S30). Interestingly, around 70% of the detected features were in 5 kbp gene flanking regions; nevertheless, only 20% of them were found in 5kbp DEG-flanking regions (Table S30). Moreover only 1% of all genes were found to be differentially methylated (Table S31), which suggests a conserved pattern of gene regulation across genotypes.

**Figure 6.**
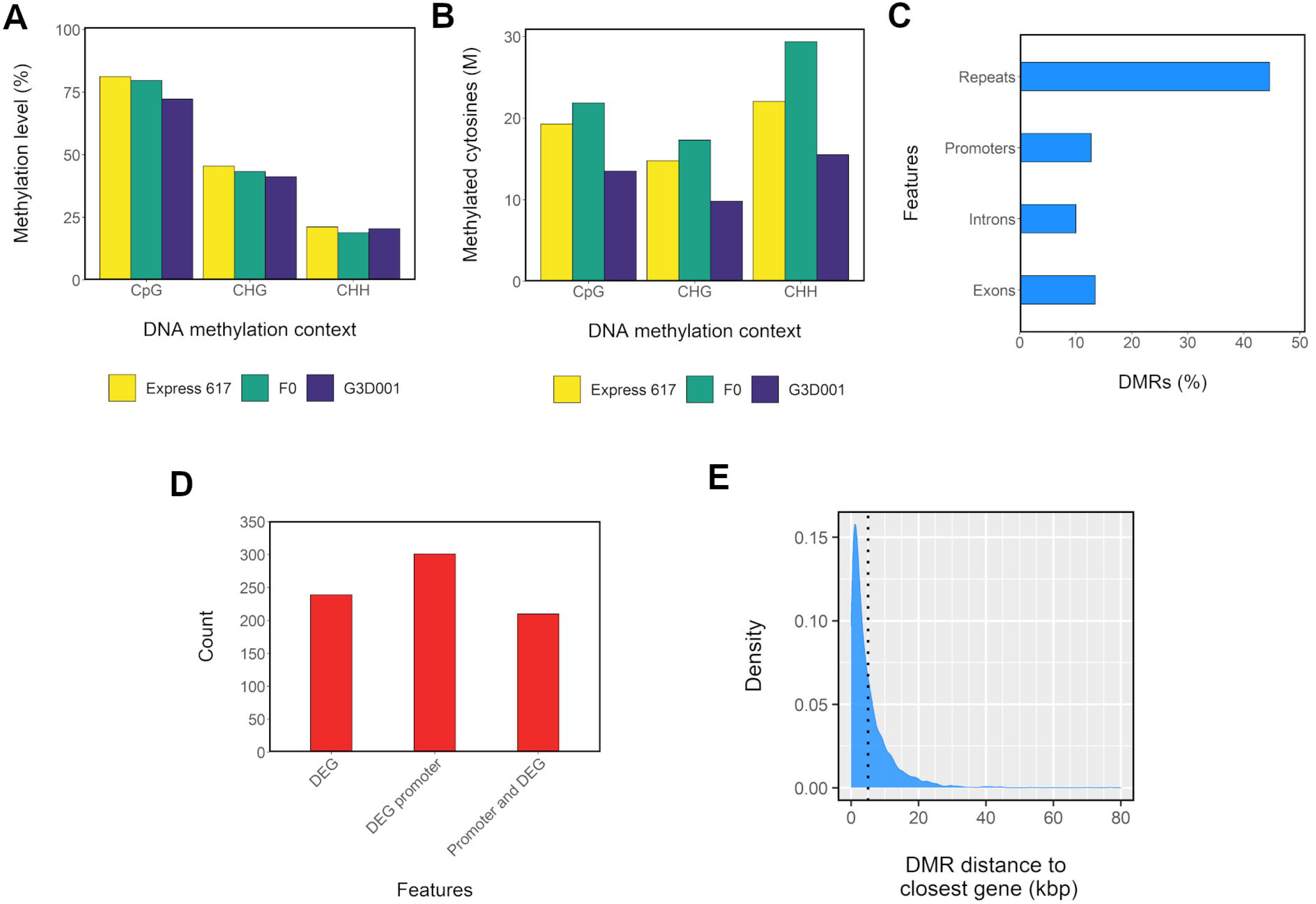
Methylation patterns in 15 days after pollination ovules from F0 and parents. **a** Methylation level per genotype and DNA methylation context. **b** Count of methylated cytosines in million (M) scale per genotype and DNA methylation context. **c** Distribution of differentially methylated regions (DMRs) across introns, exons, repeats and promoters (1 kbp upstream from gene start). **d** Distribution of methylated differential expressed genes (DEGs) and their promoters. **e** Kernel density estimation (KED)-based distribution of DMRs distance to closest gene. A dotted line is used to delimit DMRs located 5 kbp from a gene.

A total of 392 genes that were both differentially expressed and differentially methylated can be regarded as putative epialleles (Table S32). No gene ontology enrichment was found in regard to these putative epialleles. Instead, they cover diverse biological functions such as DNA transcription, carbohydrate and lipid metabolic processes and photosynthesis (Table S33). Interestingly, both the gene body and its promoter were methylated in most putative genetic epialleles (Table S34). Most DMRs were less than 5 kbp away from a gene, suggesting a potential regulatory role (Fig. 6, Figs S11-S14). Both proportional and inversely proportional relationships were detected between gene methylation and gene expression in most stage comparisons (Fig. S15-S17). However, during early seed development in the hybrid, proportional interactions with upregulation of hypomethylated genes were most prevalent (Fig. 7). Studies linking methylation and expression and evaluating epialleles have proved beneficial in detecting heterotic patterns in other crops like maize, rice and Arabidopsis (Greaves et al., 2015; Cao et al., 2022; Wang and Wang, 2022). In addition, 112635 CpG islands were detected in all assigned chromosomes in the Express 617 reference genome, with an average length of 363 bp and strong differences in frequency in centromeric regions of different chromosomes (Table S35). Although 86% of all identified CpG islands were methylated, with an average methylation level of 62%, only 1.35% of these were differentially methylated (Table S36).

**Figure 7.**
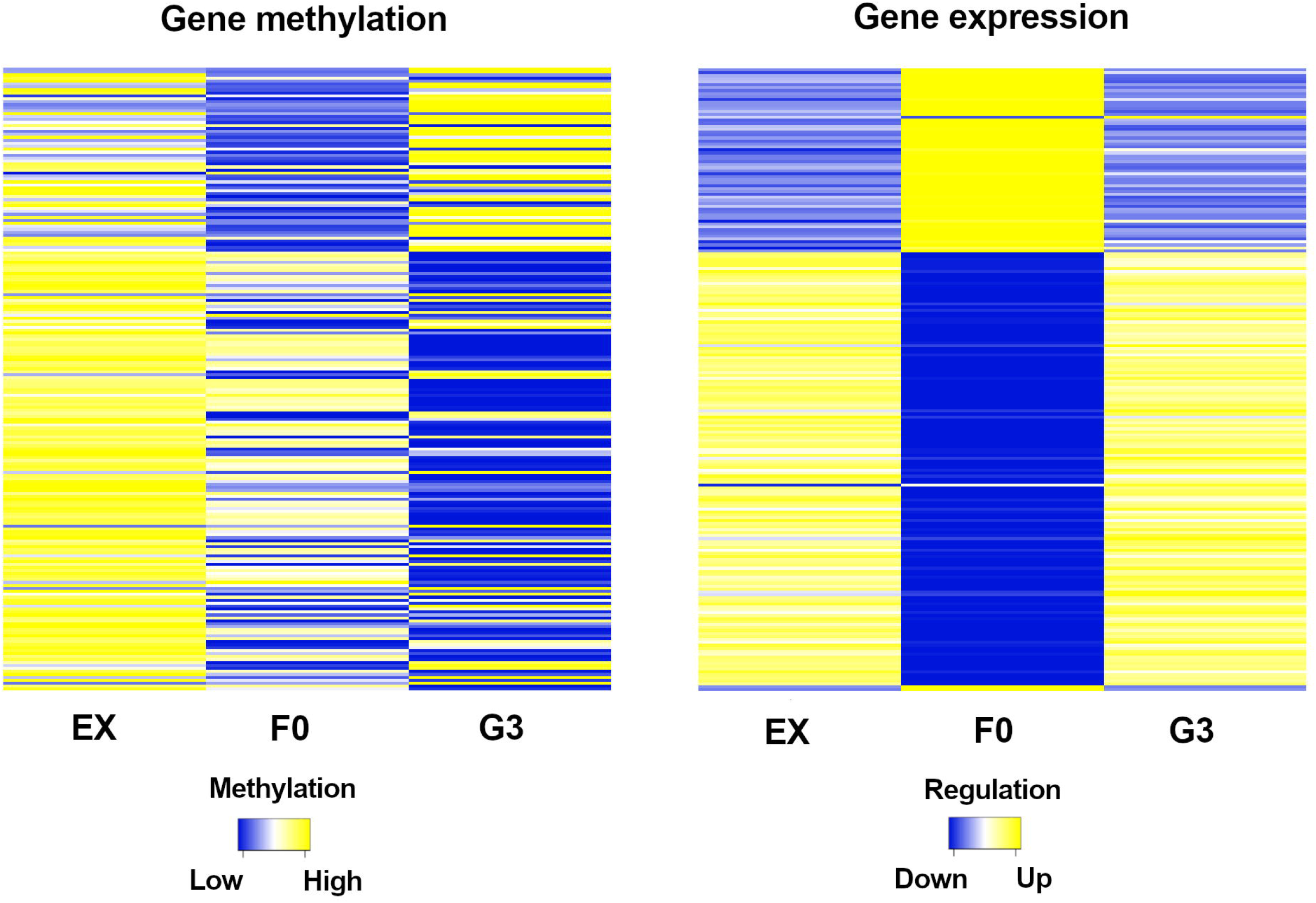
Gene expression and gene methylation in CpG and CHG contexts from 15 days after pollination ovules displaying transgressive patterns in the F0 and its parents. Genes are sorted in the same order in both heatmaps.

### Segmental and subgenome expression bias in hybrids

Segmental patterns of differential gene expression were visualised by circos plots displaying expression patterns for each chromosome, genotype and stage. The presence of putative expression clusters was assessed more precisely through a 500 kbp genome-wide binning approach where consistent DEGs patterns per segment, chromosome, genotype and stage were grouped as described in Material and Methods. Consequently, 144 differentially expressed segments across genotypes and stages were determined (Table S37). More differentially expressed segments were found in the A subgenome than the C subgenome and most segments found in F0 comparisons mimicked the expression patterns of the maternal parent Express 617 (Table S37). An example on chromosome A03 is shown in Fig. 8. Sequence reads from G3D001 pollinated ovules were employed to exclude the possibility that the observed patterns were due to genomic rearrangements (Table S38). Large scale deletions were found only in chromosome C01 in G3D001, which accounts for the low expression found on the deleted segments in that chromosome (Fig. S18-S20, Table S37). However, no large-scale rearrangements were found in chromosome A03 in G3D001 (Fig. S21), so that cannot be the reason why a segment on this chromosome showed low expression in both early (15 days) and late (30 days) seed development stages (Fig. 8 and Fig. S22). Moreover, the corresponding chromosome region in Express 617 does not appear to be duplicated, since neither the F1 nor the F0 showed a high expression pattern that could have been inherited from a large-scale duplication from the Express 617 parent (Fig. S23-24). Furthermore, no relationships were observed between methylation level, repeat density or position relative to the centromere. This suggests that specific chromosome segments may correspond more closely to maternal expression patterns than other regions. The mechanisms of such a phenomenon could be associated to parental roles during embryo development, genomic imprinting or chromatin activity and/or genome accessibility for transcription. However, more detailed investigations are necessary to validate these hypotheses.

**Figure 8.**
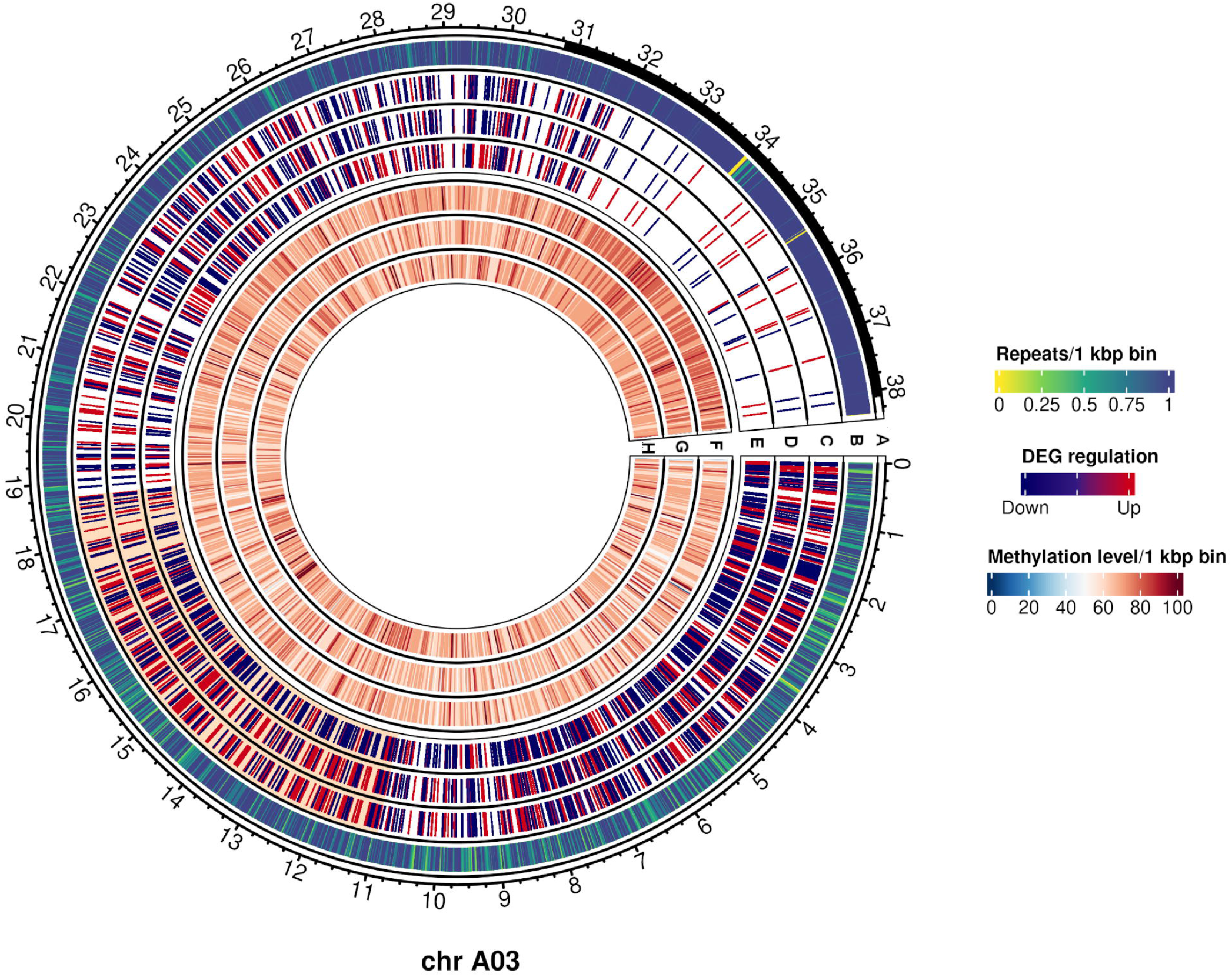
Differentially expressed genes (DEGs) and methylation levels from 15 days after pollination ovules from F0 and parents in chromosome A03. Outer to inner tracks correspond to: **a** Predicted centromere positions in black; **b** Repeat density per 1 kbp bin; **c-e** DEG regulation in **(c)** Express 617, **(d)** F0 and **(e)** G3D001; **f-h:** Methylation levels per 1 kbp bin in **(f)** Express 617, **(g)** F0 and **(h)** G3D001. A differentially expressed chromosome segment between around 11 Mbp and 18.8 Mbp is highlighted in orange in tracks c-e.

## Discussion

The results of this study demonstrate that differential expression and methylation patterns potentially associated with heterotic patterns are already detectable during seed development and seedling stages in an F1 hybrid. Most DEGs showed maternal and paternal dominances regardless of the tissue and stage, indicating that some of these differential features represent stable regulatory patterns with a general involvement in heterosis. Similar kinds of parental gene expression dominance have been reported previously in oilseed rape and cotton (Yoo et al., 2013; Wu et al., 2018; Wei et al., 2021). Li et al. (2020) demonstrated that expression dominance levels in interspecific hybrids can vary based on the sampled tissue, with stems and leaves showing more additive gene expression in allopolyploid *B. napus* compared to the expression from its diploid progenitors *B. rapa* and *B. oleracea*. Gene expression additivity was also reported by Zhang et al. (2021a) as a main pattern of expression dominance levels in excised pod sections in crosses between *Raphanus sativus* (RR, 2n =18) and *B. oleracea*, whereas seeds and pods from the homozygous diploids displayed predominantly paternal dominance. The diversity of sampled tissues, species and genotypes in the previous studies and ours could account for the contrasting expression dominance levels observed. The segmental patterns of differential gene expression on a chromosome scale which we observed reflect reports in *B. napus* by Lloyd et al. (2018) and He et al. (2017), who found that large scale rearrangements induced large segmental expression differences. Interestingly, however, not all differentially expressed chromosome segments analyzed in our study were associated with large scale rearrangements. Although the underlying reasons for this observation require further elucidation, this may indicate that chromosome-level patterns of chromatin rearrangement and transcription accessibility may be involved.

Additionally, pollinated F0 ovules that develop later into F1 seeds and plants showed a higher similarity to the maternal genotype Express 617 in terms of gene expression, small RNA expression and methylation, As noted by Jahnke et al. (2010), this might be due to the triploid nature of the endosperm, which arises from the union of a duplicated maternal gamete and a paternal gamete via double-fertilization. Seeds are composed of a seed coat, embryo and endosperm, with the proportions of the three components varying depending on species and developmental age. Transcriptomic profiling of different seed tissues using laser microdissection has been employed to characterize the transcriptomic profiles during seed formation in *A. thaliana* and *B. napus* (Kirkbride et al., 2019; Ziegler et al., 2019; Khan et al., 2022), and could provide further insight into heterotic patterns during F1 seed development, however this technically demanding task was outside of the scope of the present study.

Subgenomic expression bias has been reported earlier in *Brassica* species (Bird et al., 2018; Bird et al., 2021a; Zhang et al., 2021b). Hence we also investigated expression bias of differentially expressed up- and downregulated genes on a subgenomic basis in each genotype and stage. Although we did not detect any subgenome bias in gene expression, on the whole more genes were differentially upregulated in G3D001 than in Express 617 (Table S15, Figures S1-S5). The observed genotype-specific bias is potentially a result of different genomic, transcriptomic and epigenomic factors. Firstly, genomic rearrangements gene copy number variations and other structural variants are known to affect various traits in *B. napus* and other polyploid plants (Schiessl et al., 2017; Vollrath et al., 2021; Makhoul et al., 2022) and could have led to potential biases in expression patterns. Transcriptomic aspects such as gene isoforms, gene network interactions and allele expression bias might also be involved in favoring the up- or down-regulation from a certain genotype or haplotype (Fan et al., 2020; Schiessl et al., 2020; Golicz et al., 2021). Lastly, epigenomic factors like parental gamete methylation mechanisms, genomic imprinting or differences between parental *cis-trans* regulating factors, miRNA isoforms (isomiRs) and TE families and densities could all result in potential genotype-biased or haplotype-biased expression (Jain et al., 2018; Go and Civetta, 2020; Gill et al., 2021).

Around 12-18% of features shared in at least 2-3 stages followed maternal dominant patterns, highlighting the potential relevance of maternal effects on transcriptomic and epigenomic regulation of early development in this hybrid. Furthermore, less than 3% of all expression and methylation features had the same expression and methylation patterns across all stages, suggesting that the role of those features might be more essential throughout seed and early seedling development. Our results in regard to GO enrichment of differentially regulated features also underlined the role of heterotic patterns in driving key biological functions involved in photosynthesis, growth and development. Heterosis has been associated to a combination of similar functions like photosynthetic activity and cell division, which already help to enhance performance during early developmental stages (Liu et al., 2021). Here, DEGs involved in reproduction and meiotic functions already showed transgressive upregulated expression in ovules from selfed-F1 plants at 15 days after pollination. Information on these genes and their expression patterns could be of potential interest for approaches to use transcriptomic data for prediction of hybrid performance.

Interestingly, we observed differentially expressed miRNAs in early and late seed development among miRNA families which are normally involved in plant growth and development (Plotnikova et al., 2019; Dong et al., 2022; Verma et al., 2022). For example, miR172 regulates not only the flowering time pathway, but also embryo development by controlling *APETALA 2* (*AP2*) and *AP2-like* genes such as *TOE2* (Boutilier et al., 2002; Shivaraj et al., 2018; Nowak et al., 2022). miR165/166 families control leaf adaxial/abaxial development and embryogenesis by targeting the class III homeodomain leucine zipper (HD-ZIP III) transcription factor gene family which includes the *REV*, *PHV* and *PHB* (Wang et al. 2007, Tang et al. 2012). Both *PHB* and *PHV* have been described to indirectly regulate *LEC2*, a gene that promotes embryo formation in Arabidopsis and seed size and seed lipid biosynthesis in *B. napus* (Braybrook et al., 2006; Tang et al., 2012; Wójcik et al., 2017; Miller et al., 2019). As noted by Dong et al. (2022), miR169 targets *C-REPEAT BINDING FACTOR* (*CBF*) and *NUCLEAR FACTOR YA* (*NF-YA*) genes. During late seed development in *B. napus*, we found that miR169 targets *EMB2016*, a member of the EMB gene family critical for embryo development (Tzafrir et al., 2004; Růžička et al., 2017; Meinke, 2020), while *EMB2204* was targeted by miR3629. mir3629 was first reported in *Vitis vinifera* cv. Pinot Noir by Pantaleo et al. (2010) and has since been reported in *Camellia azalea*, in response to chilling in *Prunus persica* and in disease susceptibility in *V. vinifera* cv. Bosco and *V. vinifera* cv. Chardonnay (Barakat et al., 2012; Pantaleo et al., 2016; Yin et al., 2016; Snyman et al., 2017). mir9410 has been detected in *B. oleracea* and *B. rapa* (Lukasik et al., 2013; Zhang et al., 2018), yet no clear function information for mir9410 exists for Brassica species. In our study, miR9410 targeted a *filamentation temperature sensitive protein H 1* (*FtsH7*) gene copy encoding a protease that in turns degrades D1 protein in photosystem II. *FtsH* genes have been reported in tomato, sorghum, Arabidopsis and *B. napus* (Xu et al., 2021; Yi et al., 2022).

The study identified multiple differential miRNA sequences and their putative targets. Further validation of targets associated with DE-miRNAs can potentially be achieved through degradome sequencing (German et al., 2008), precise isomiRs classification (Morin et al., 2008; Sablok et al., 2015; Yang et al., 2019), target knock-out experiments (Jain et al., 2018; Wei et al., 2018; Li et al., 2021) or gene co-expression networks (Schiessl et al., 2020) in order to delimit their role in seed and embryo formation in *B. napus*.

Overall, the number of methylated cytosines in the CHH context was higher in all genotypes compared to other contexts; nonetheless, methylation levels were higher in CpG and CHG contexts, as observed previously in multiple plants species (e.g. Niederhuth et al., 2016; Bartels et al., 2018). Methylation is generally associated with gene downregulation through transcription inhibition. Nevertheless, hypermethylation and hypomethylation were also linked with up- and downregulation, respectively. Proportional gene hypermethylation and upregulation was observed in mice and human cells (Arechederra et al., 2018; Rauluseviciute et al., 2020); however, no mechanisms explaining gene activation through hypermethylation are fully known so far; thus, further research would elucidate the interactions between methylation and gene regulation, particularly in relation to heterozygosity and heterosis.

In our study we evaluated methylation during seed development because parental asymmetric methylation and genomic imprinting occurs mostly at that stage in flowering plants (Batista and Köhler, 2020). DNA hypomethylation of the female gamete and paternal gamete hypermethylation has been reported in many flowering plants, including Arabidopsis, rice and maize (Gehring et al., 2009; Zemach et al., 2010; Zhang et al., 2014b). Interestingly, we observed contrasting patterns of maternal hypermethylation and paternal hypomethylation in the F0. Similar parental methylation trends were also observed in the F1 despite allele segregation. Such patterns were also reported by Liu et al. (2018) in *B. napus* and by Grover et al. (2020) in *B.rapa*. As a possible explanation for maternal hypermethylation, Grover et al. (2020) proposed that a high expression of so-called “siren” siRNAs in the seed coat could trigger maternal DNA methylation during seed development. The molecular mechanisms and effects of genomic imprinting, where an allele follows a parental expression pattern due to inherited epigenomic modifications, is restricted mostly to the endosperm rather than the embryo in flowering plants, and has been extensively discussed by Weigel and Colot (2012) and Batista and Köhler (2020), respectively. The role of imprinted genes has been linked to chromatin modification, hormone biosynthesis, nutrient transfer, endosperm proliferation and seed size regulation (reviewed by Jiang and Köhler 2012, Batista and Köhler 2020). Furthermore, Rong et al. (2021) reported the enrichment of transposable elements located in 5 kbp flanking regions of imprinted genes in *B. napus*. Cao et al. (2022) analyzed imprinted genes in six backcrossing generations of maize as well as in three selfing generations derived from the 6^th^ backcross. They proposed that the divergence between TEs derived from 24-nt siRNAs in the parental maize genomes might have led to transgenerational inheritance of imprinted genes. Putative imprinted genes were also found in the seedling and seed development stages in our study. Epigenetic changes have been reported as relevant heterotic factors which are influenced by allele diversity, parental effects and environmental conditions (Botet and Keurentjes, 2020). Epigenomic parental effects are more likely to occur during seed formation, when gametes fuse to form a zygote, given that this stage is marked by epigenomic features involving siRNA, DNA methylation, imprinting and chromatin activity. Because these shape the epigenomic and transcriptomic landscape of the zygote and, potentially, its future development as a seedling, hybrids can potentially benefit strongly from a heterotic advantage imparted by these features in these very early developmental stages.

Most frequently, differentially methylated transposable elements corresponded to abundant Copia and Gypsy families. Transposable elements are key factors in speciation and subgenome expression patterns (Bird et al., 2018; Bottani et al., 2018; Bird et al., 2021a) and are known for their high variability across plant species (Novák et al., 2020; Mhiri et al., 2022). TEs can also alter the epigenetic landscape in relation to hybrid fitness (Serrato-Capuchina and Matute, 2018). Therefore, detailed assessment of transposable element densities and compositions between hybrids and their parents could be beneficial. In addition, siRNAs are known to mediate silencing of transposable elements via the RNA-directed DNA methylation (RdDM) pathway. At the same time, TEs are a source of sRNAs, including siRNAs, that could potentially silence TEs through a post-transcriptional gene silencing (PTGS) process (Matzke and Mosher 2014; Gill et al. 2021). As described by Rong et al. (2021), most differentially methylated TEs were found in 5 kbp gene flanking regions. Because most TEs, DMRs and DE-siRNAS converged into regions directly flanking genes, whereas there was no abundance in regions flanking DEGs, gene regulation in the hybrid appears to be associated with conservation of key genetic functions by reducing the number of DMRs, DE-siRNAs and differentially methylated TEs in the proximity of DEGs. This could be one of the most interesting and significant results of the present study. Additionally, most CpG islands were not differentially methylated between genotypes, indicating a putative role in regulating gene expression, as reported in rice and Arabidopsis (Ashikawa, 2001).

The sequencing data gathered from single tissues in the present study allowed an integrated view of gene expression, small RNA interactions and genomic methylation during early developmental stages in an oilseed rape hybrid developed from two distant genotypes. Coding and non-coding features which were differentially expressed or methylated in this study provide new insight into early expression of heterosis in oilseed rape seeds and seedlings from a molecular viewpoint. The extent of these features in an allopolyploid model crop like *B. napus* also have potential implications in other polyploid crops where heterosis still remains to be exploited, such as wheat and potatoes (Steeg et al., 2022). Patterns of expression and methylation dominance levels could also contribute a new level of understand regarding allele-specific gene expression (Fan et al., 2020; Sands et al., 2021), isoform expression (Vitting-Seerup and Sandelin, 2019; Yao et al., 2020; Golicz et al., 2021), gene fusion and dosage (Mahmoud et al., 2019; Serin Harmanci et al., 2020; Bird et al., 2021b) as well as non-germline omics variations among F1 plants and populations (Higgins et al., 2018; Cortijo et al., 2019; Orantes-Bonilla et al., 2022; Quezada-Martinez et al., 2022). Their role in heterotic gene expression patterns is ultimately also of interest for transcriptome-based genomic selection or hybrid performance prediction (e.g. Frisch et al., 2010). Defining the roles of differentially expression regulatory features in early developmental stages of hybrids could be used to enhance expression-based prediction model (Seifert et al., 2018b; Zrimec et al., 2020; Cheng et al., 2021; Hu et al., 2021b; Knoch et al., 2021). Altogether our findings highlight transcriptomics and epigenomic differences between early developmental stages in F1 and F0 in terms of methylation level as well as in gene and small RNA expression. The contribution of differential coding and non-coding features to early hybrid seed formation is of key interest for hybrid breeding and deserves further evaluation using more diverse genotypes, heterotic pools and in different species. Future developments in sequencing and bioinformatics will also aid in elucidating the role and interactions among transcriptomic and epigenomic features at higher resolution, helping to expand current knowledge and applications of heterosis in polyploid crops.

## Supplementary Information

Supplementary figures and tables are available in the online article version.

## Acknowledgements

The authors acknowledge support for bioinformatic resources from the BMBF-funded de.NBI Cloud within the German Network for Bioinformatics Infrastructure (de.NBI) and the Bioinformatics Core Facility at JLU. The experimental design was created with BioRender.com.

## Author Contributions statement

RJS and JZ conceived and supervised the study. MOB drafted the manuscript and designed the bioinformatic analyses. HW conducted bioinformatic studies, generated crosses, extracted and sampled pollinated ovules for sequencing and contributed to data analysis. DH carried and supervised the field experiment. WL contributed to experimental trials. HTL performed genome synteny analyses and contributed to wholegenome bisulfite and mRNA analyses. AAG contributed to analyses of transcriptomic and epigenomic features. All authors read and approved the manuscript.

## Data availability

mRNA, sRNA and WGBS libraries and fragment count datasets generated in this study are found in the GEO data repository under accession GSE202610. G3D001 genomic reads from self-pollinated ovules are found in NCBI Bioproject PRJNA850551.

