## Supplementary Figures for "Transgressive and parental dominant gene expression and cytosine methylation during seed development in *Brassica napus* hybrids"

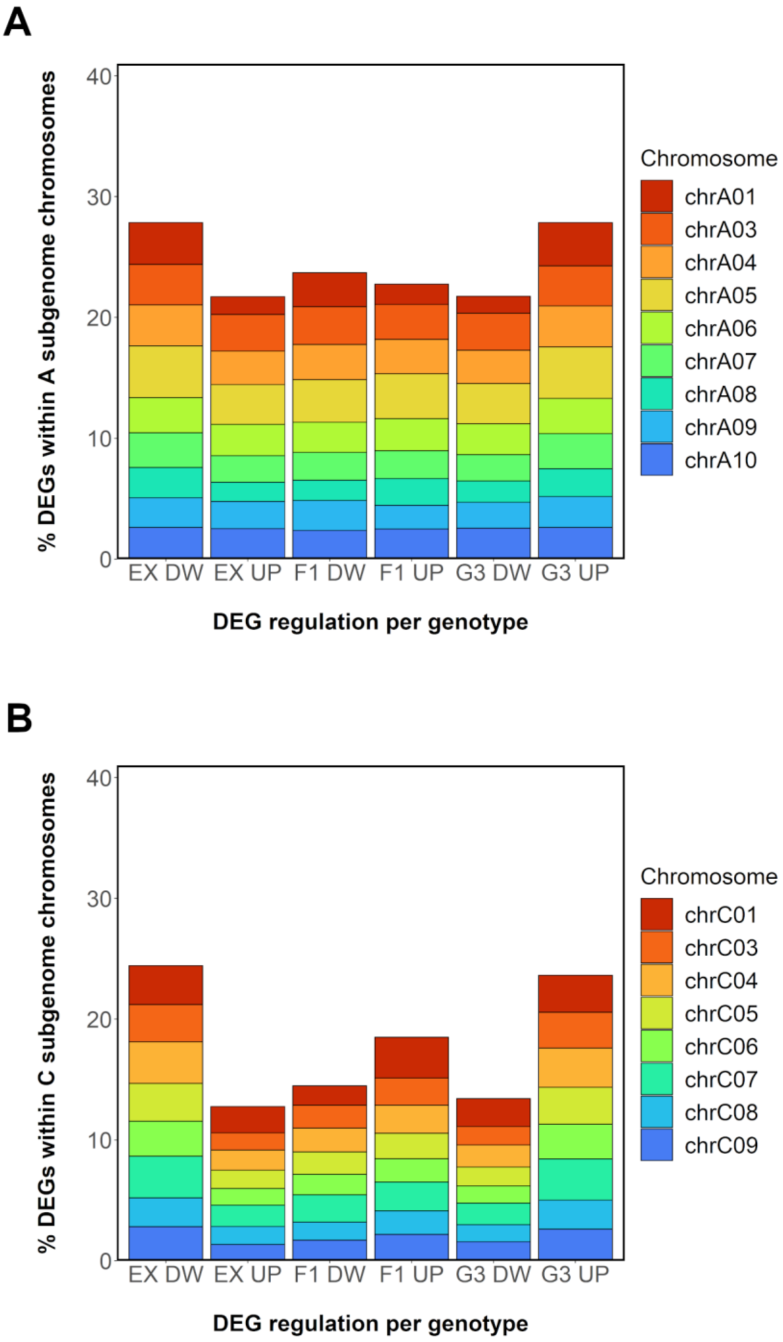


**Supplementary Figure 1** Percentage of expressed upregulated and downregulated differentially expressed genes (DEGs) per subgenome and genotype during seedling stage (BBCH16). **a** Percentages of DEGs in respect to all genes per chromosome in A subgenome. **b** Percentages of DEGs in respect to all genes per chromosome in C subgenome. DEGs in genotypes (EX: Express 617, G3: G3D001 and F1) are classified either as upregulated (UP) or downregulated (DW). Percentages are calculated based on the number of DEGs observed in each subgenome. Chromosomes A02 and C02 were excluded to discard analysis bias due to whole A02 chromosome duplication and whole chromosome C02 deletion observed in G3D001 (Orantes-Bonilla et al., 2022).

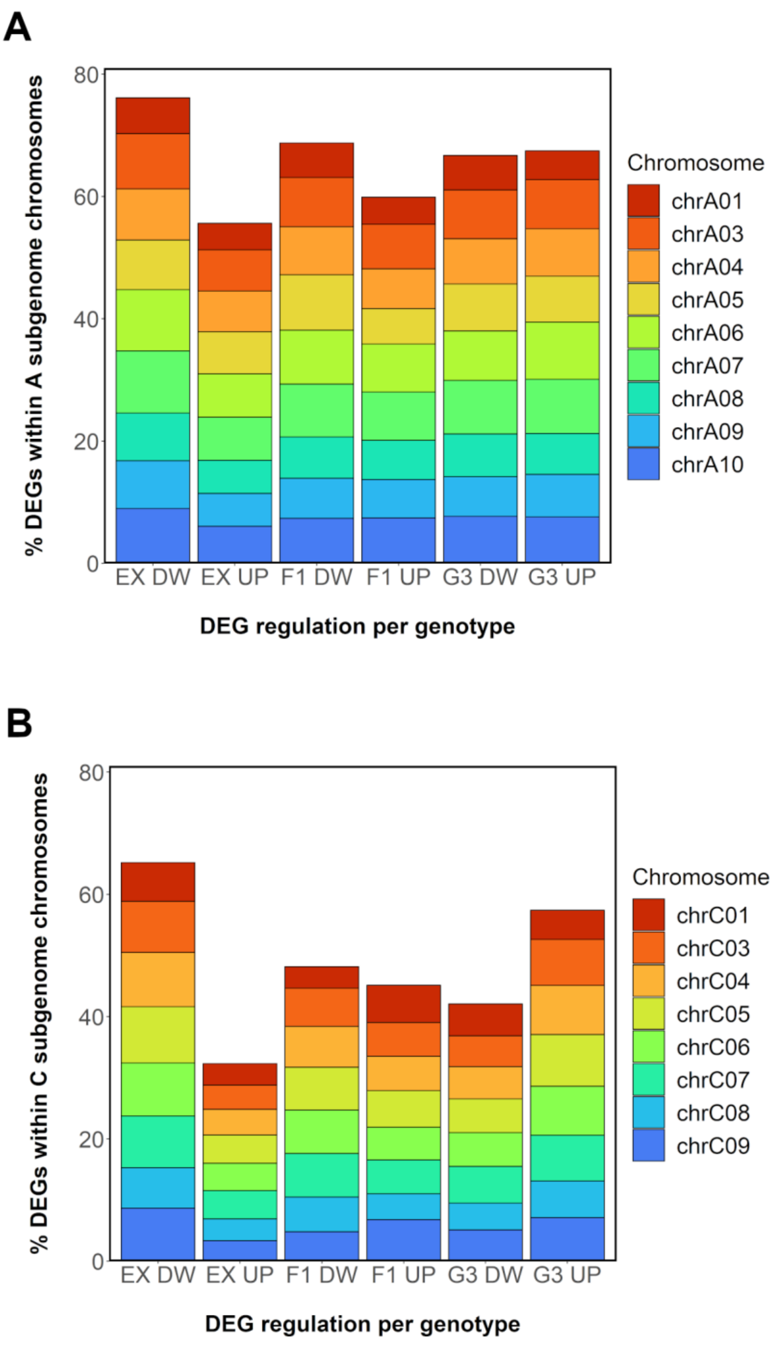


**Supplementary Figure 2** Percentage of expressed upregulated and downregulated differentially expressed genes (DEGs) per subgenome and genotype in 15 days after pollination ovules in F1 and parents. **a** Percentages of DEGs in respect to all genes per chromosome in A subgenome. **b** Percentages of DEGs in respect to all genes per chromosome in C subgenome. DEGs in genotypes (EX: Express 617, G3: G3D001 and F1) are classified either as upregulated (UP) or downregulated (DW). Percentages are calculated based on the number of DEGs observed in each subgenome. Chromosomes A02 and C02 were excluded to discard analysis bias due to whole A02 chromosome duplication and whole chromosome C02 deletion observed in G3D001 (Orantes-Bonilla et al., 2022).

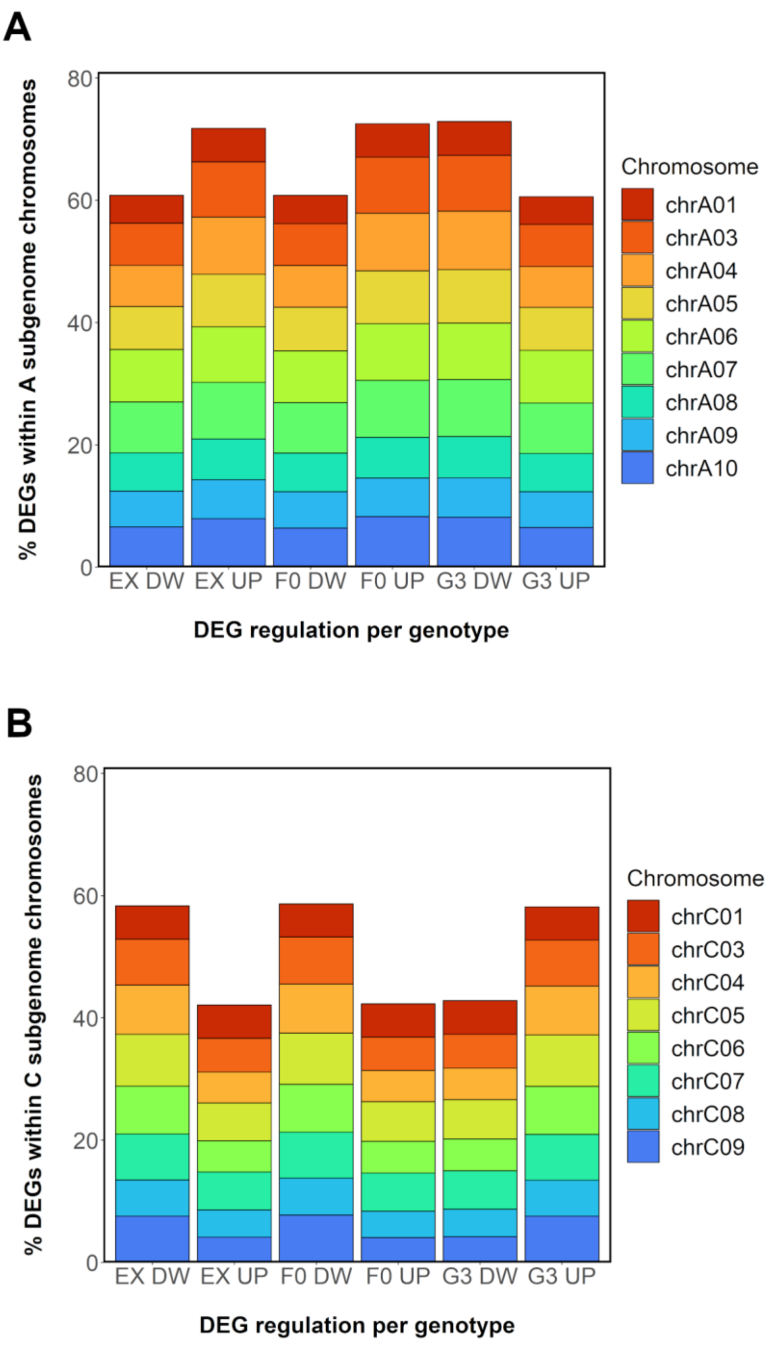


**Supplementary Figure 3** Percentage of expressed upregulated and downregulated differentially expressed genes (DEGs) per subgenome and genotype in 15 days after pollination ovules in F0 and parents. **a** Percentages of DEGs in respect to all genes per chromosome in A subgenome. **b** Percentages of DEGs in respect to all genes per chromosome in C subgenome. DEGs in genotypes (EX: Express 617, G3: G3D001 and F0) are classified either as upregulated (UP) or downregulated (DW). Percentages are calculated based on the number of DEGs observed in each subgenome. Chromosomes A02 and C02 were excluded to discard analysis bias due to whole A02 chromosome duplication and whole chromosome C02 deletion observed in G3D001 (Orantes-Bonilla et al., 2022).

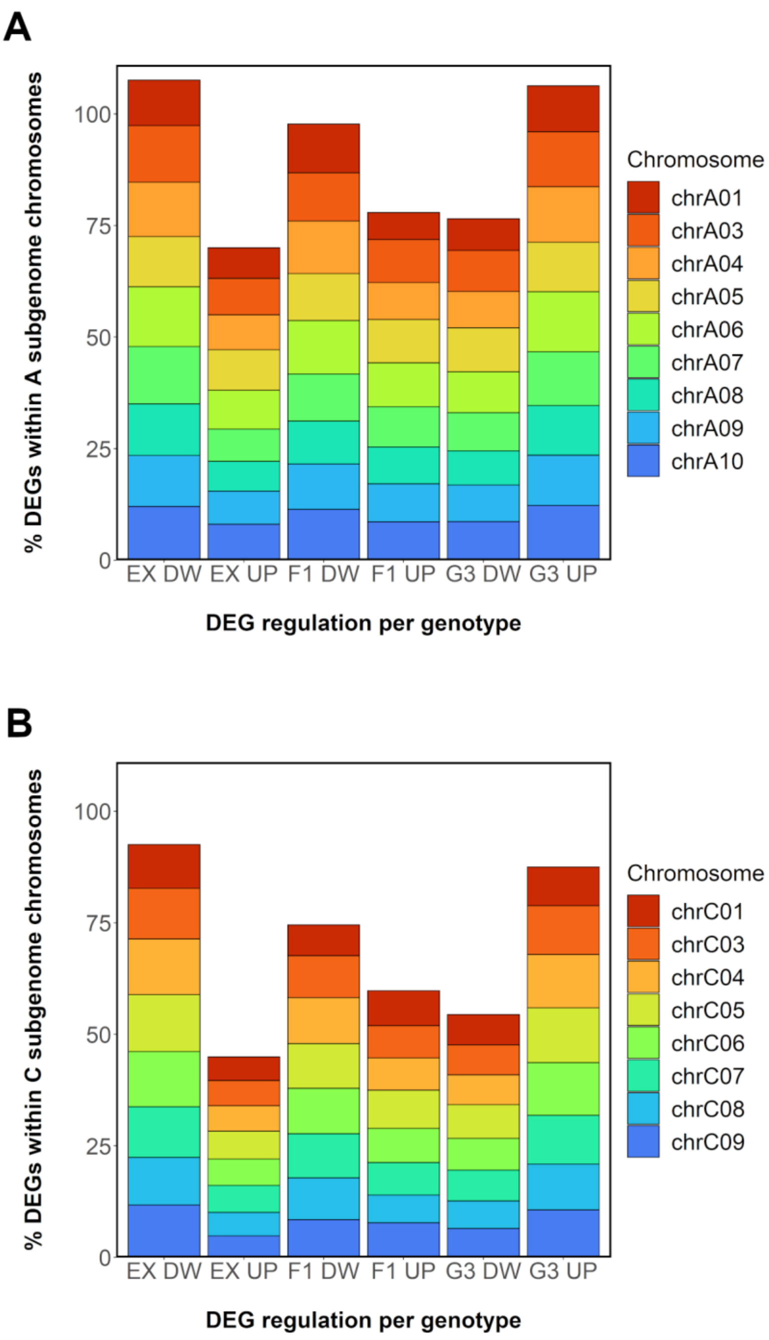


**Supplementary Figure 4** Percentage of expressed upregulated and downregulated differentially expressed genes (DEGs) per subgenome and genotype in 30 days after pollination ovules in F1 and parents. **a** Percentages of DEGs in respect to all genes per chromosome in A subgenome. **b** Percentages of DEGs in respect to all genes per chromosome in C subgenome. DEGs in genotypes (EX: Express 617, G3: G3D001 and F1) are classified either as upregulated (UP) or downregulated (DW). Percentages are calculated based on the number of DEGs observed in each subgenome. Chromosomes A02 and C02 were excluded to discard analysis bias due to whole A02 chromosome duplication and whole chromosome C02 deletion observed in G3D001 (Orantes-Bonilla et al., 2022).

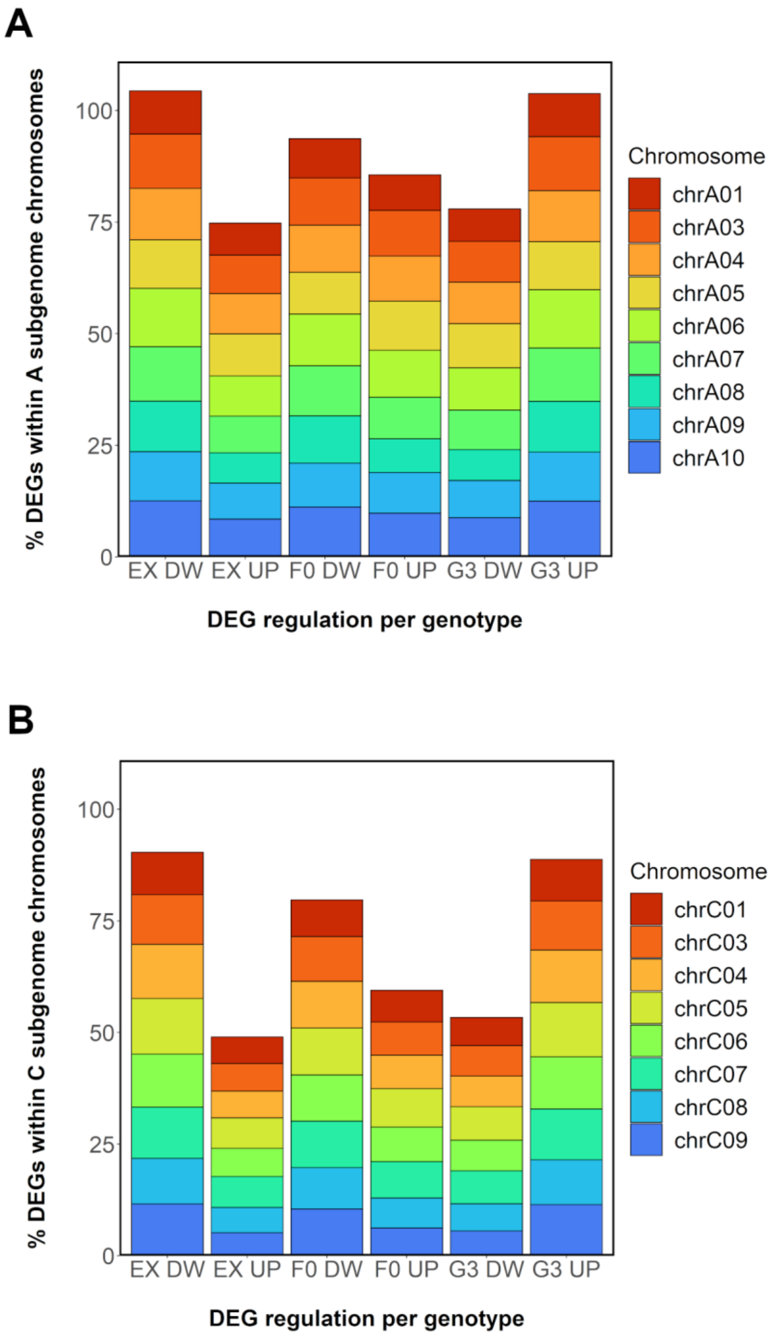


**Supplementary Figure 5** Percentage of expressed upregulated and downregulated differentially expressed genes (DEGs) per subgenome and genotype in 30 days after pollination ovules in F0 and parents. **a** Percentages of DEGs in respect to all genes per chromosome in A subgenome. **b** Percentages of DEGs in respect to all genes per chromosome in C subgenome. DEGs in genotypes (EX: Express 617, G3: G3D001 and F0) are classified either as upregulated (UP) or downregulated (DW). Percentages are calculated based on the number of DEGs observed in each subgenome. Chromosomes A02 and C02 were excluded to discard analysis bias due to whole A02 chromosome duplication and whole chromosome C02 deletion observed in G3D001 (Orantes-Bonilla et al., 2022).

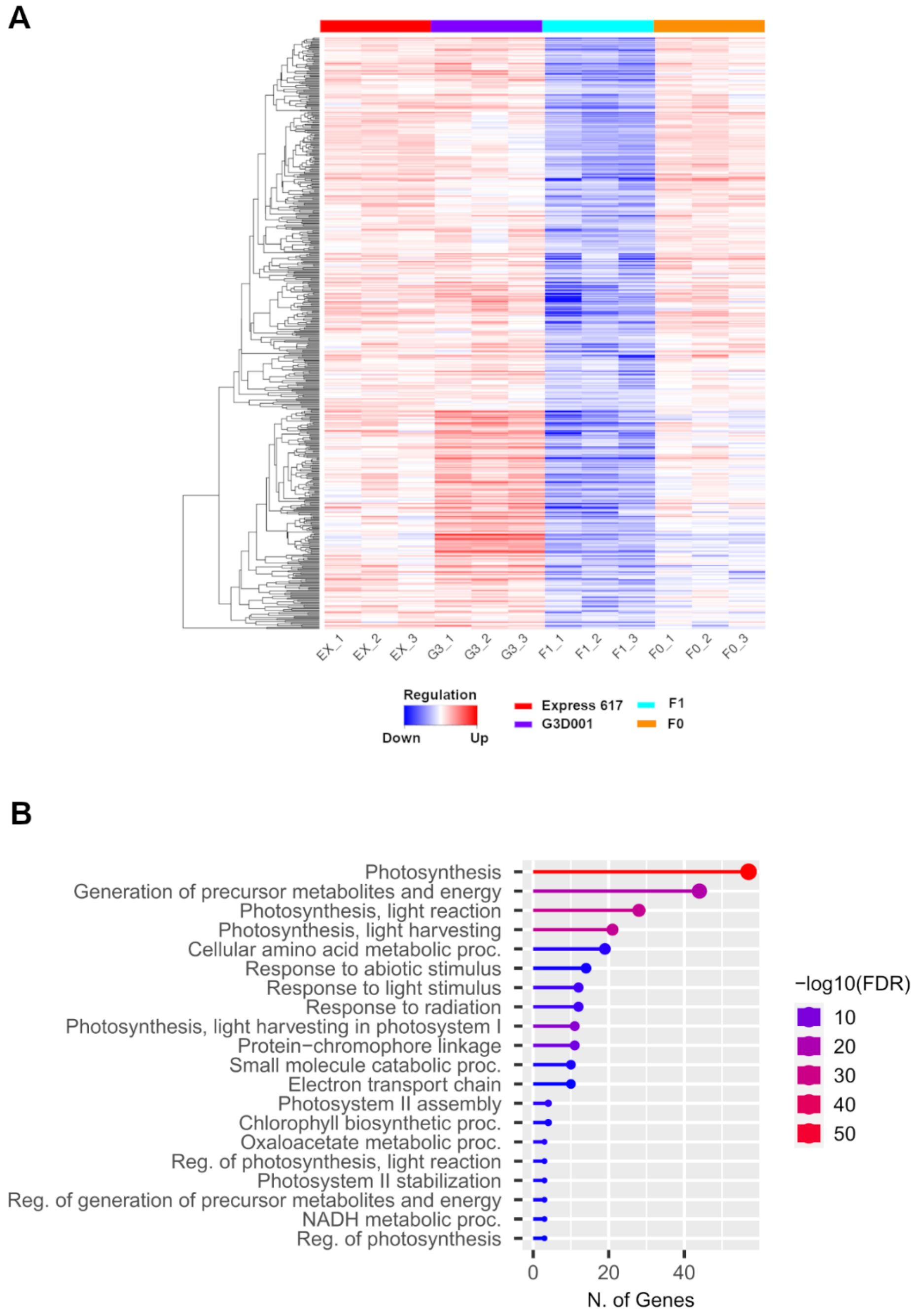


**Supplementary Figure 6** (**a**) Gene expression heatmap and (**b**) gene ontology (GO) enrichment of biological processes from 15 days after pollination ovules with transgressive downregulation patterns in the F1.

A

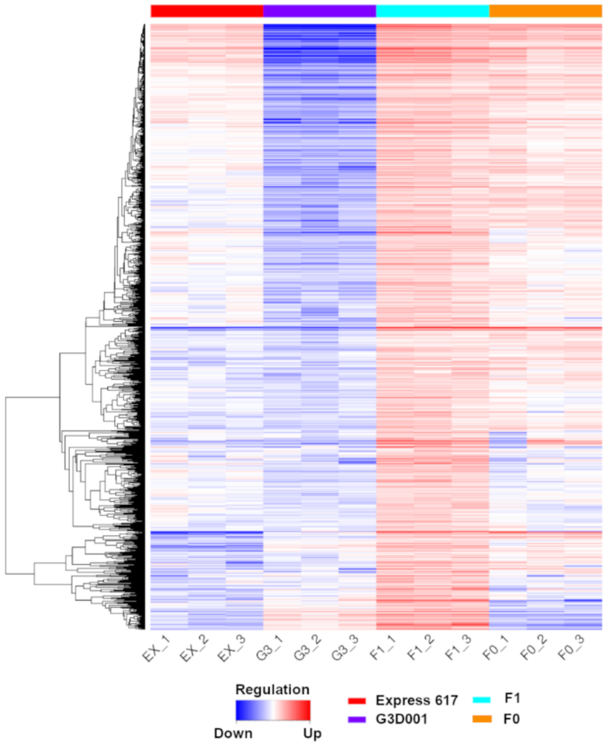


B

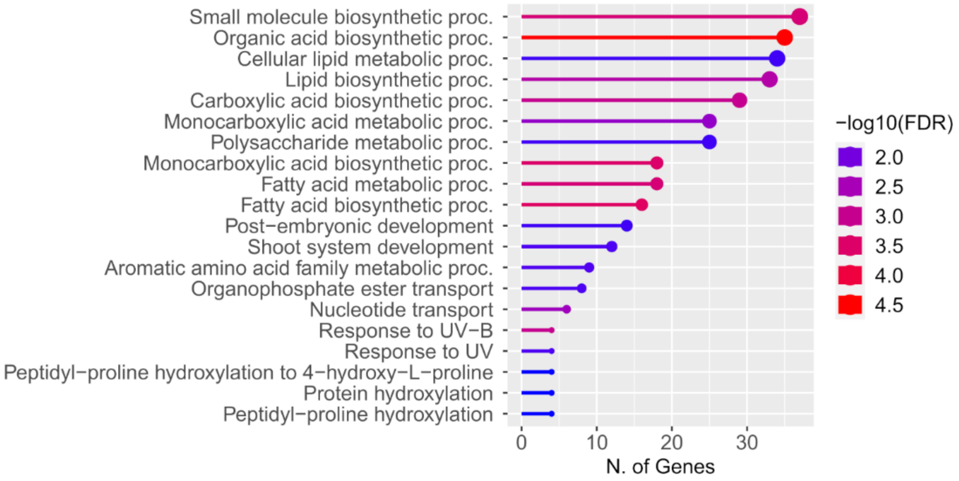


**Supplementary Figure 7** (**a**) Gene expression heatmap and (**b**) gene ontology (GO) enrichment of biological processes from 30 days after pollination ovules with transgressive upregulation patterns in the F1**.**

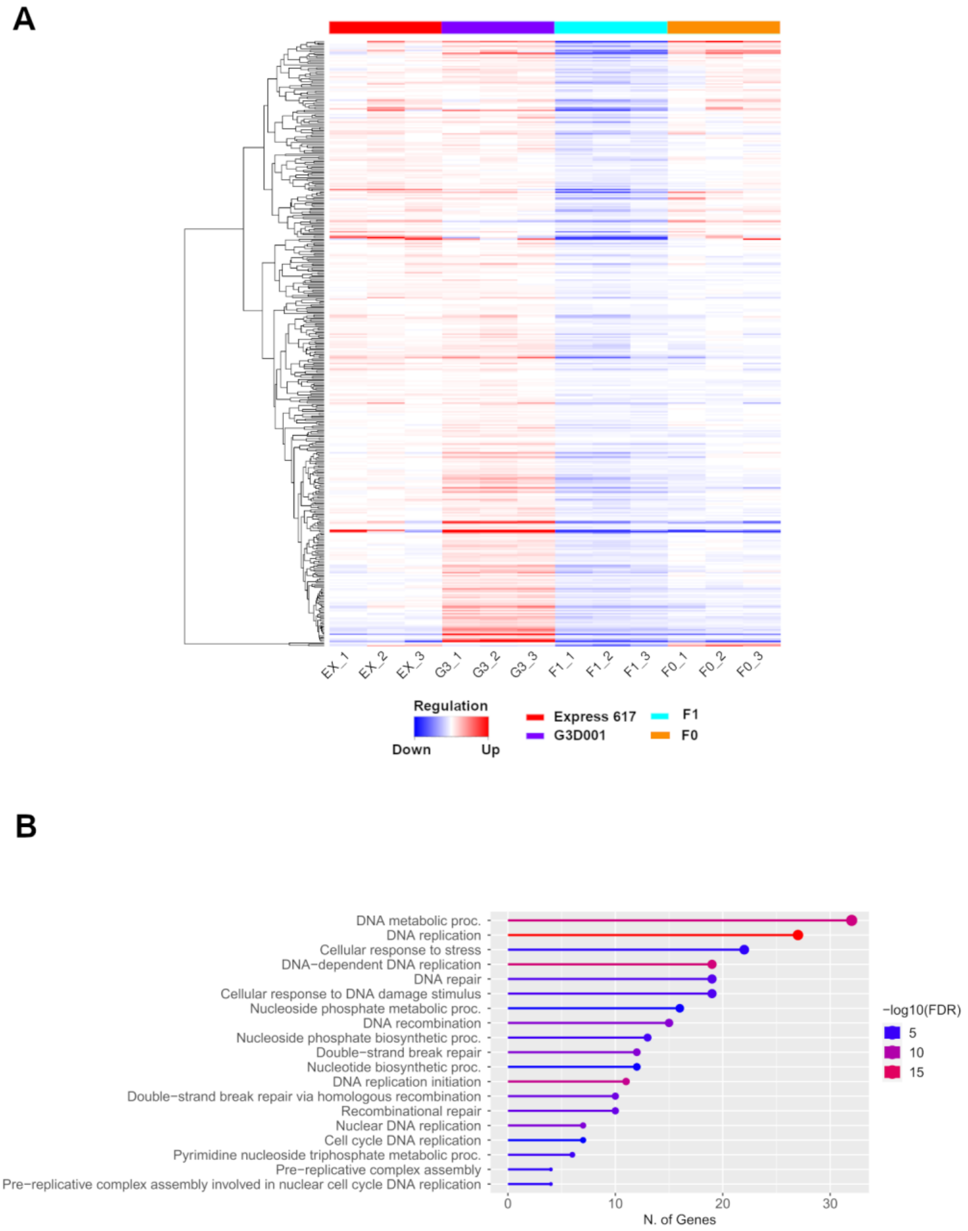


**Supplementary Figure 8** (**a**) Gene expression heatmap and (**b**) gene ontology (GO) enrichment of biological processes from 30 days after pollination ovules with transgressive downregulation patterns in the F1.

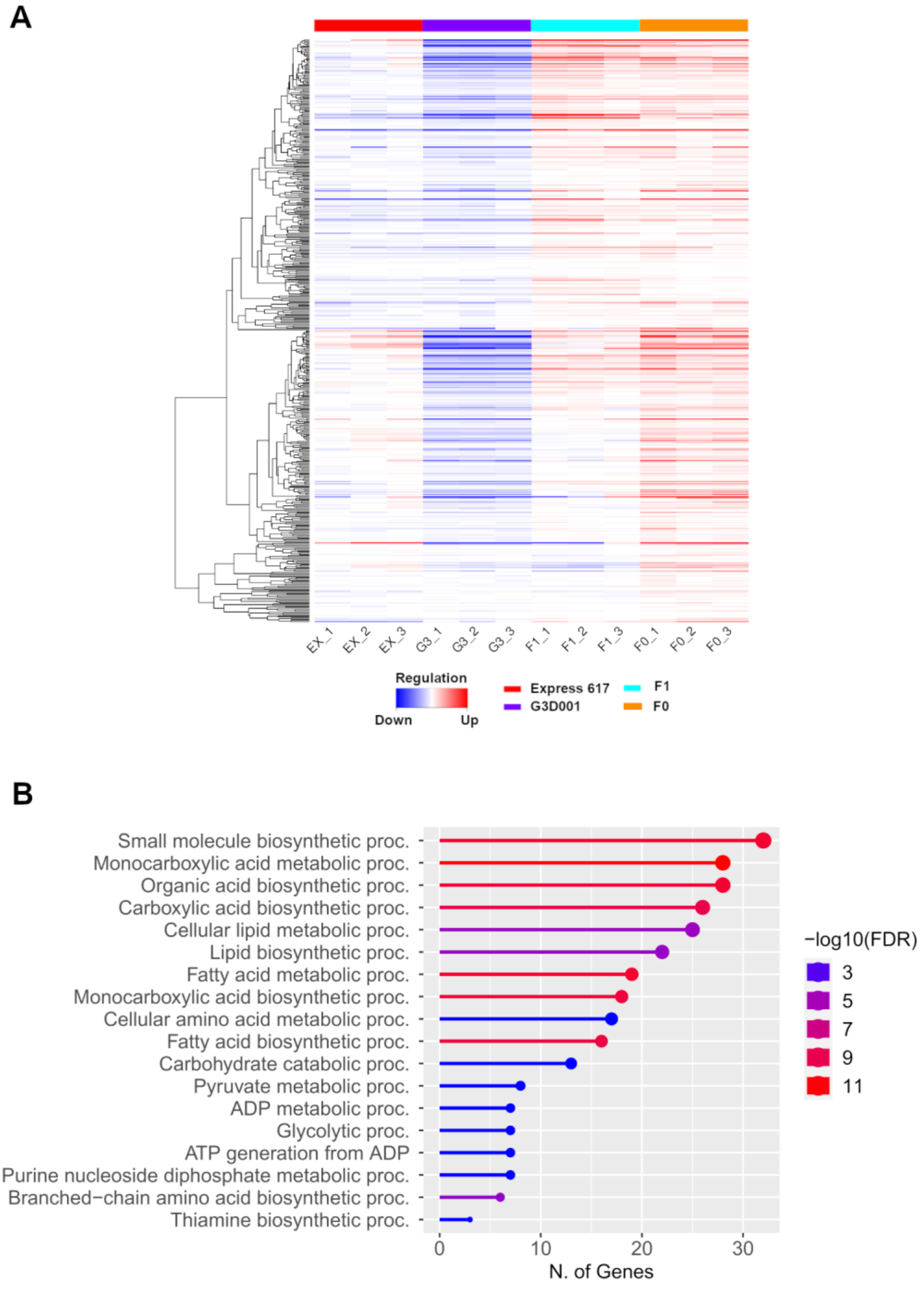


**Supplementary Figure 9** (**a**) Gene expression heatmap and (**b**) gene ontology (GO) enrichment of biological processes from 30 days after pollination ovules with transgressive upregulation patterns in the F0.

A

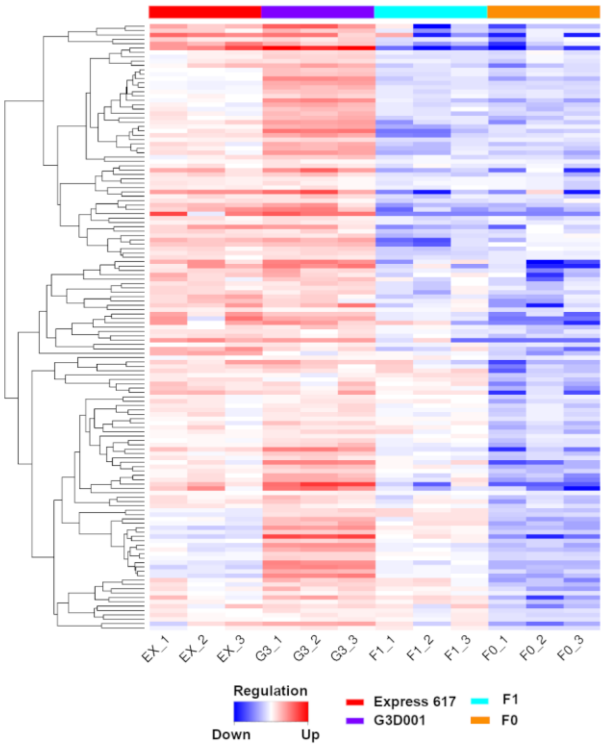


B

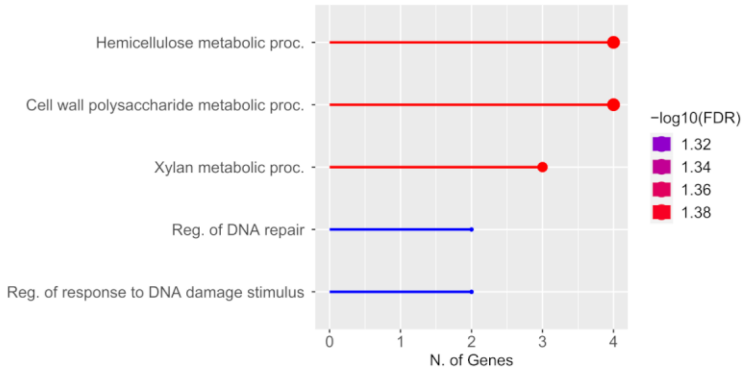


**Supplementary Figure 10** (**a**) Gene expression heatmap and (**b**) gene ontology (GO) enrichment of biological processes from 30 days after pollination ovules with transgressive upregulation patterns in the F0.

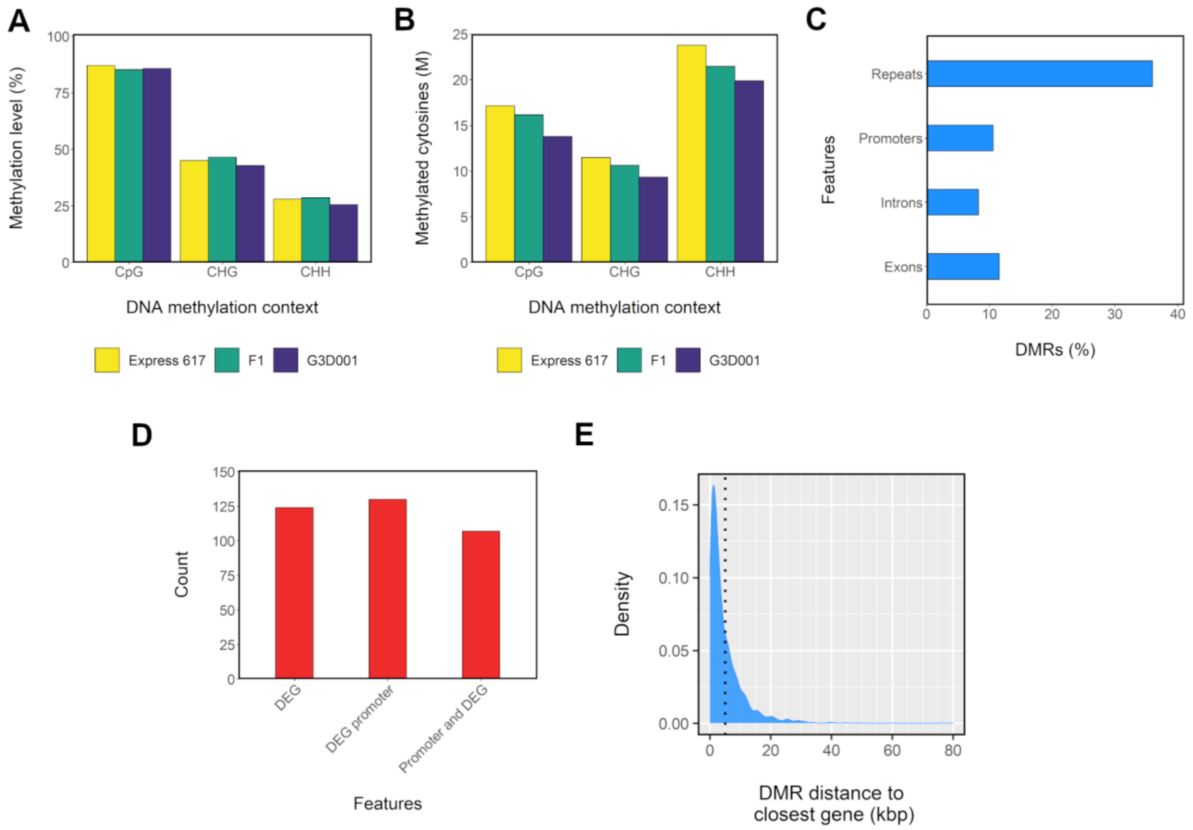


**Supplementary Figure 11** Methylation patterns in seedling stage (BBCH16) from F1 and parents. **a** Methylation level per genotype and DNA methylation context. **b** Count of methylated cytosines in million (M) scale per genotype and DNA methylation context. **c** Distribution of differentially methylated regions (DMRs) across introns, exons, repeats and promoters (1 kbp upstream from gene start). **d** Distribution of methylated differential expressed genes (DEGs) and their promoters. **e** Kernel density estimation (KED)-based distribution of DMRs distance to closest gene. A dotted line is used to delimit DMRs located 5 kbp from a gene.

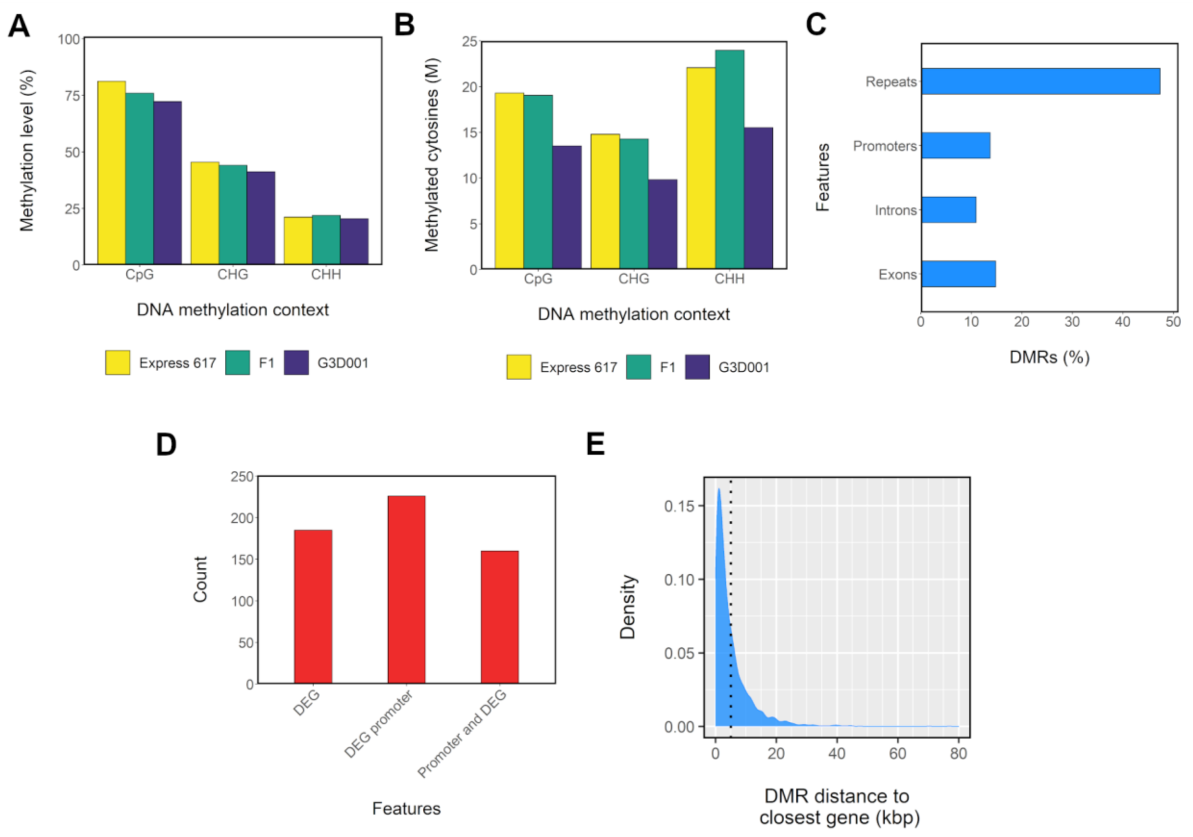


**Supplementary Figure 12** Methylation patterns in 15 days after pollination ovules from F1 and parents. **a** Methylation level per genotype and DNA methylation context. **b** Count of methylated cytosines in million (M) scale per genotype and DNA methylation context. **c** Distribution of differentially methylated regions (DMRs) across introns, exons, repeats and promoters (1 kbp upstream from gene start). **d** Distribution of methylated differential expressed genes (DEGs) and their promoters. **e** Kernel density estimation (KED)-based distribution of DMRs distance to closest gene. A dotted line is used to delimit DMRs located 5 kbp from a gene.

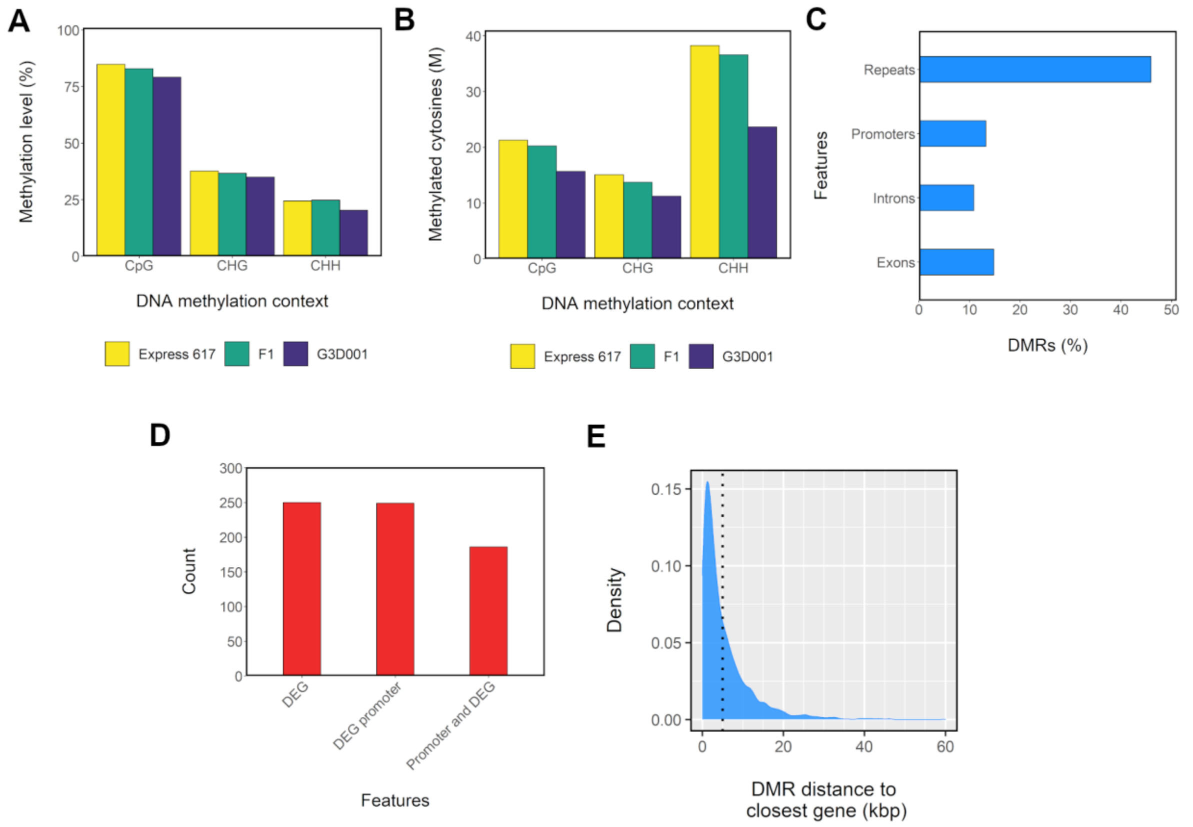


**Supplementary Figure 13** Methylation patterns in 30 days after pollination ovules from F1 and parents. **a** Methylation level per genotype and DNA methylation context. **b** Count of methylated cytosines in million (M) scale per genotype and DNA methylation context. **c** Distribution of differentially methylated regions (DMRs) across introns, exons, repeats and promoters (1 kbp upstream from gene start). **d** Distribution of methylated differential expressed genes (DEGs) and their promoters. **e** Kernel density estimation (KED)-based distribution of DMRs distance to closest gene. A dotted line is used to delimit DMRs located 5 kbp from a gene.

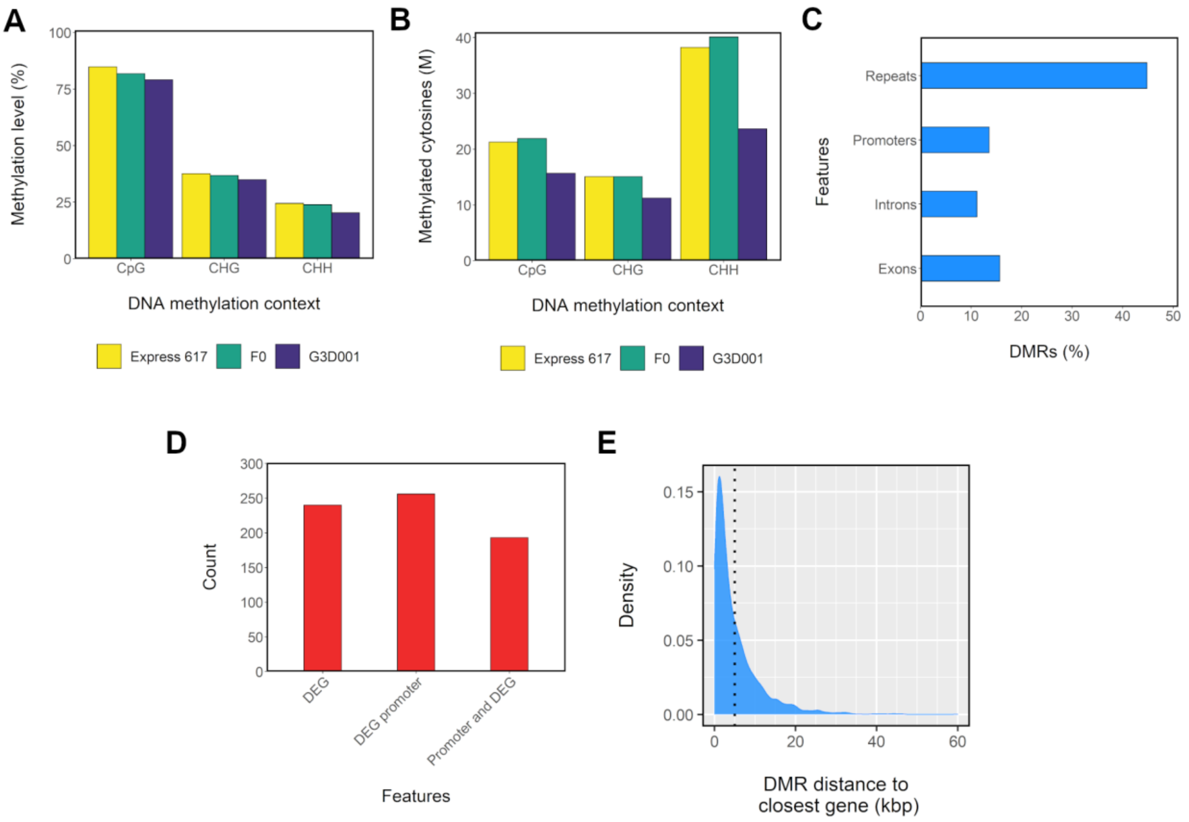


**Supplementary Figure 14** Methylation patterns in 30 days after pollination ovules from F0 and parents. **a** Methylation level per genotype and DNA methylation context. **b** Count of methylated cytosines in million (M) scale per genotype and DNA methylation context. **c** Distribution of differentially methylated regions (DMRs) across introns, exons, repeats and promoters (1 kbp upstream from gene start). **d** Distribution of methylated differential expressed genes (DEGs) and their promoters. **e** Kernel density estimation (KED)-based distribution of DMRs distance to closest gene. A dotted line is used to delimit DMRs located 5 kbp from a gene.

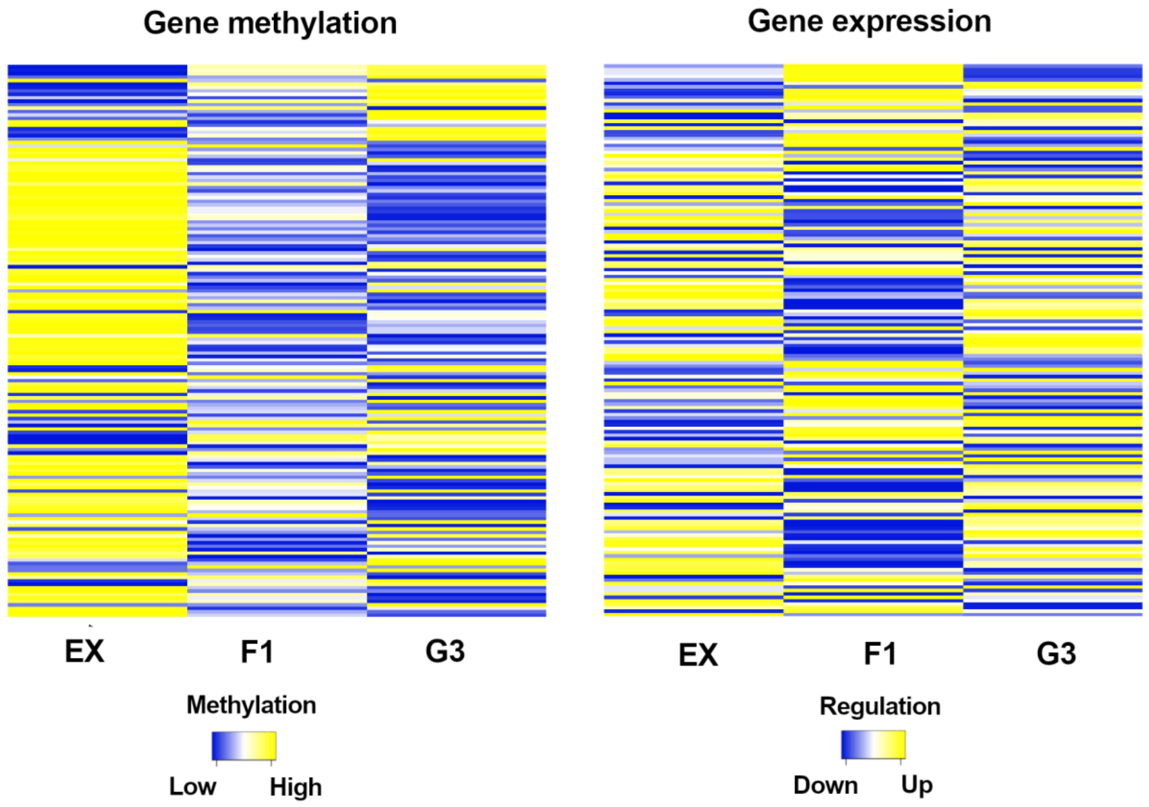


**Supplementary Figure 15** Gene expression and gene methylation in CpG and CHG contexts from 15 days after pollination ovules displaying transgressive patterns in the F1 and its parents. Genes are sorted in the same order in both heatmaps.

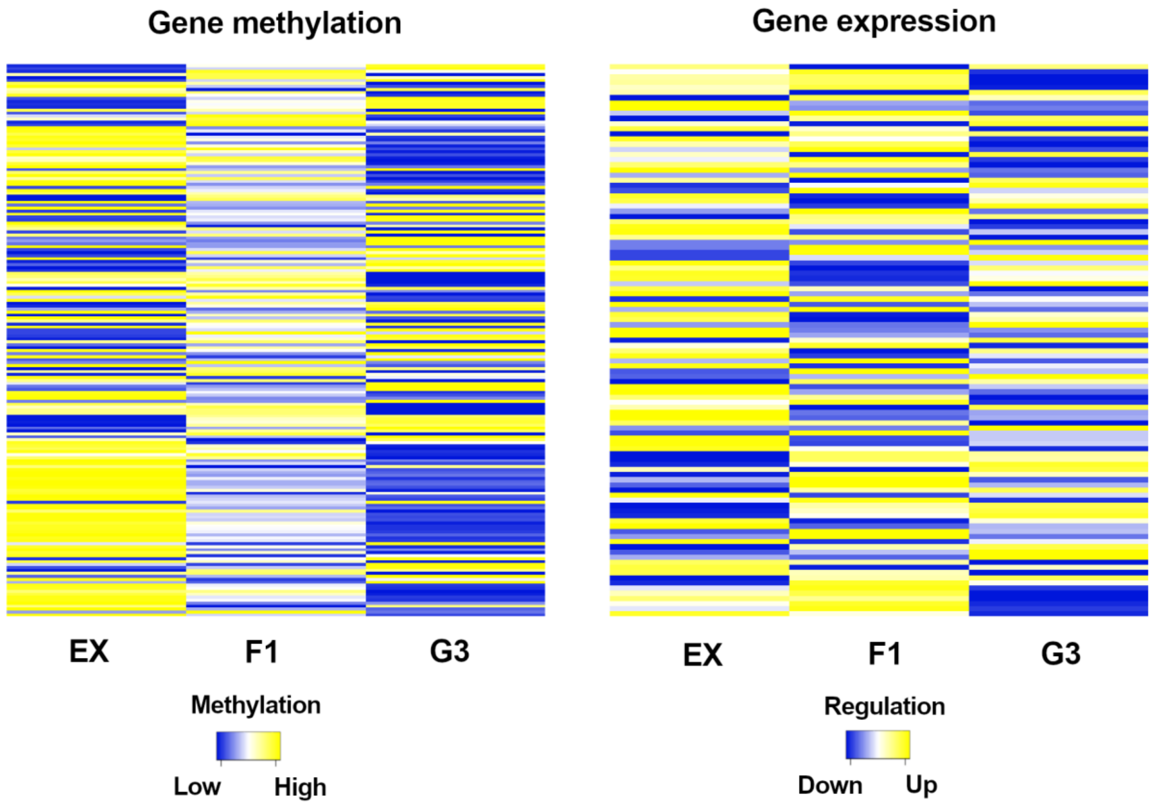


**Supplementary Figure 16** Gene expression and gene methylation in CpG and CHG contexts from 30 days after pollination ovules displaying transgressive patterns in the F1 and its parents. Genes are sorted in the same order in both heatmaps.

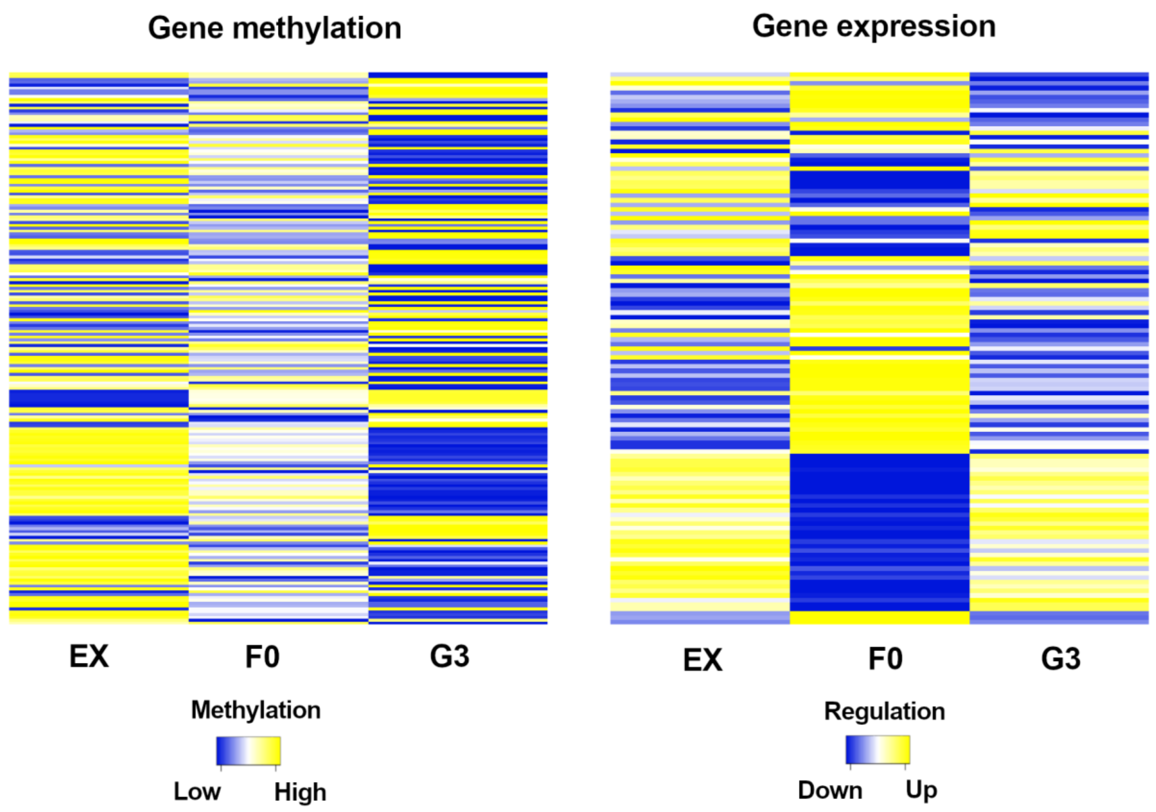


**Supplementary Figure 17** Gene expression and gene methylation in CpG and CHG contexts from 30 days after pollination ovules displaying transgressive patterns in the F0 and its parents. Genes are sorted in the same order in both heatmaps.

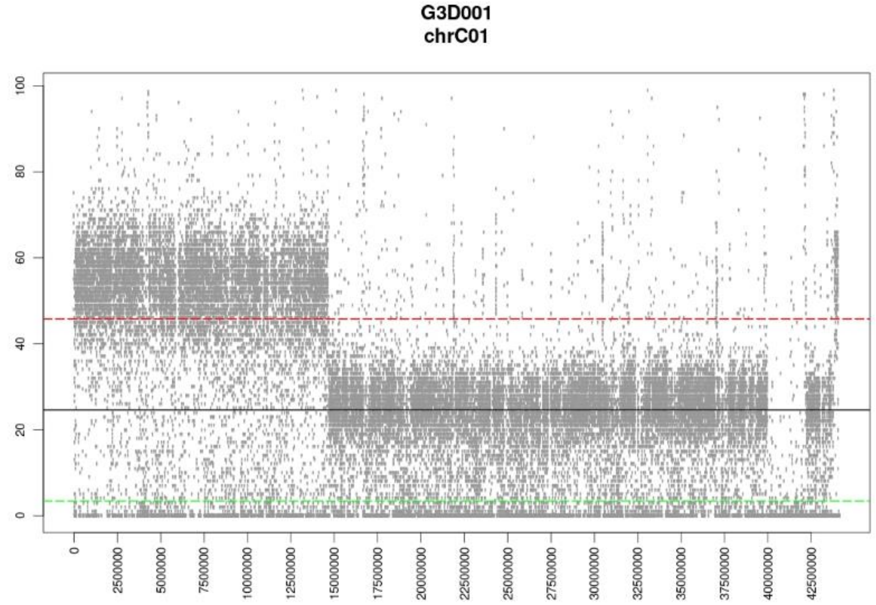


**Supplementary Figure 18** Coverage of chromosome C01 from G3D001 self-pollinated ovule based on the Express 617 reference. Duplications are shown above the red line and deletions below the green line

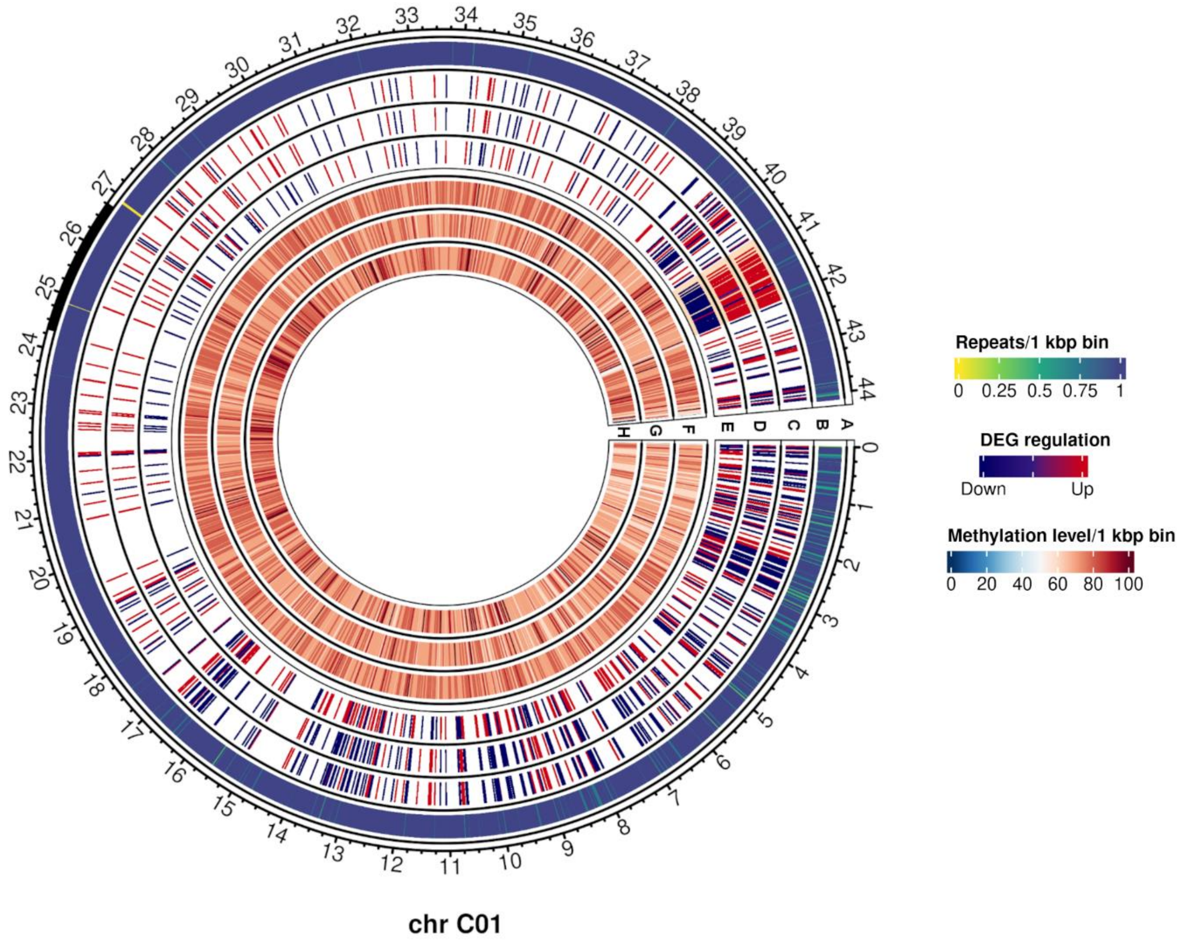


**Supplementary Figure 19** Differentially expressed genes (DEGs) and methylation levels from 15 days after pollination ovules from F0 and parents in chromosome C01. Outer to inner tracks correspond to: **a** Predicted centromere positions in black; **b** Repeat density per 1 kbp bin; **c-e** DEG regulation in **(c)** Express 617, **(d)** F0 and **(e)** G3D001; **f-h:** Methylation levels per 1 kbp bin in **(f)** Express 617, **(g)** F0 and **(h)** G3D001. A differentially expressed chromosome segment between around 40.75 Mbp and 42 Mbp is highlighted in orange in tracks c-e.

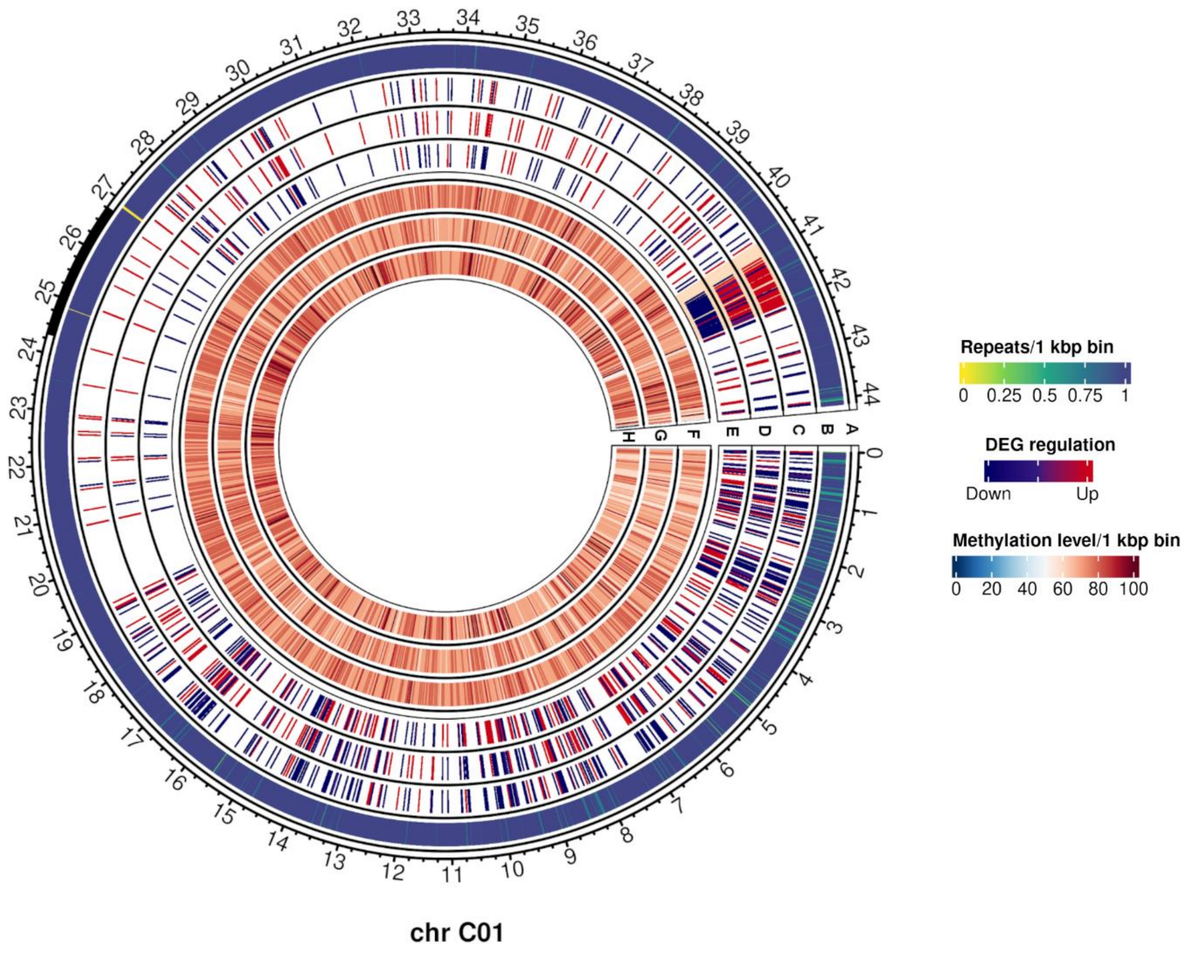


**Supplementary Figure 20** Differentially expressed genes (DEGs) and methylation levels from 15 days after pollination ovules from F1 and parents in chromosome C01. Outer to inner tracks correspond to: **a** Predicted centromere positions in black; **b** Repeat density per 1 kbp bin; **c-e** DEG regulation in **(c)** Express 617, **(d)** F1 and **(e)** G3D001; **f-h:** Methylation levels per 1 kbp bin in **(f)** Express 617, **(g)** F1 and **(h)** G3D001. A differentially expressed chromosome segment between around 40.75 Mbp and 42 Mbp is highlighted in orange in tracks c-e.

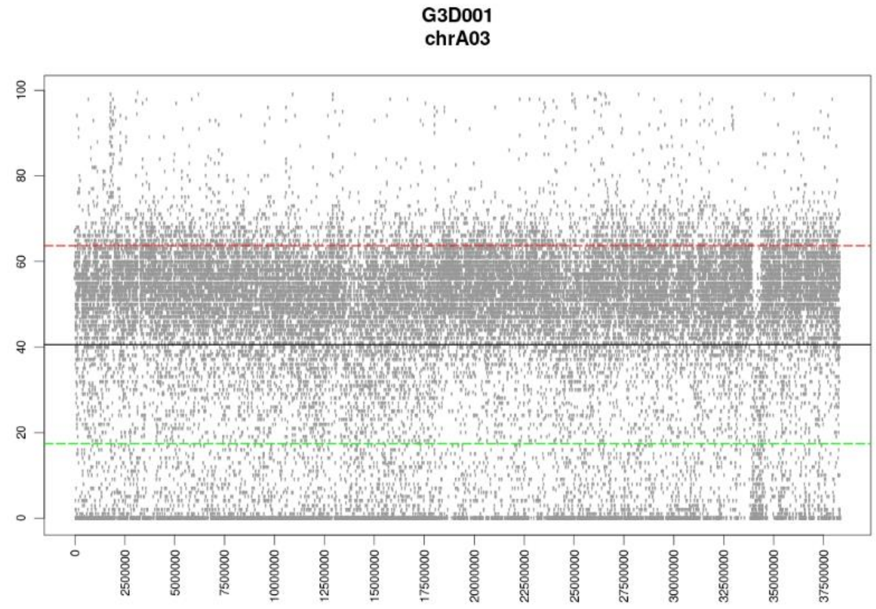


**Supplementary Figure 21** Coverage of chromosome A03 from G3D001 self-pollinated ovule based on the Express 617 reference. Duplications are shown above the red line and deletions below the green line

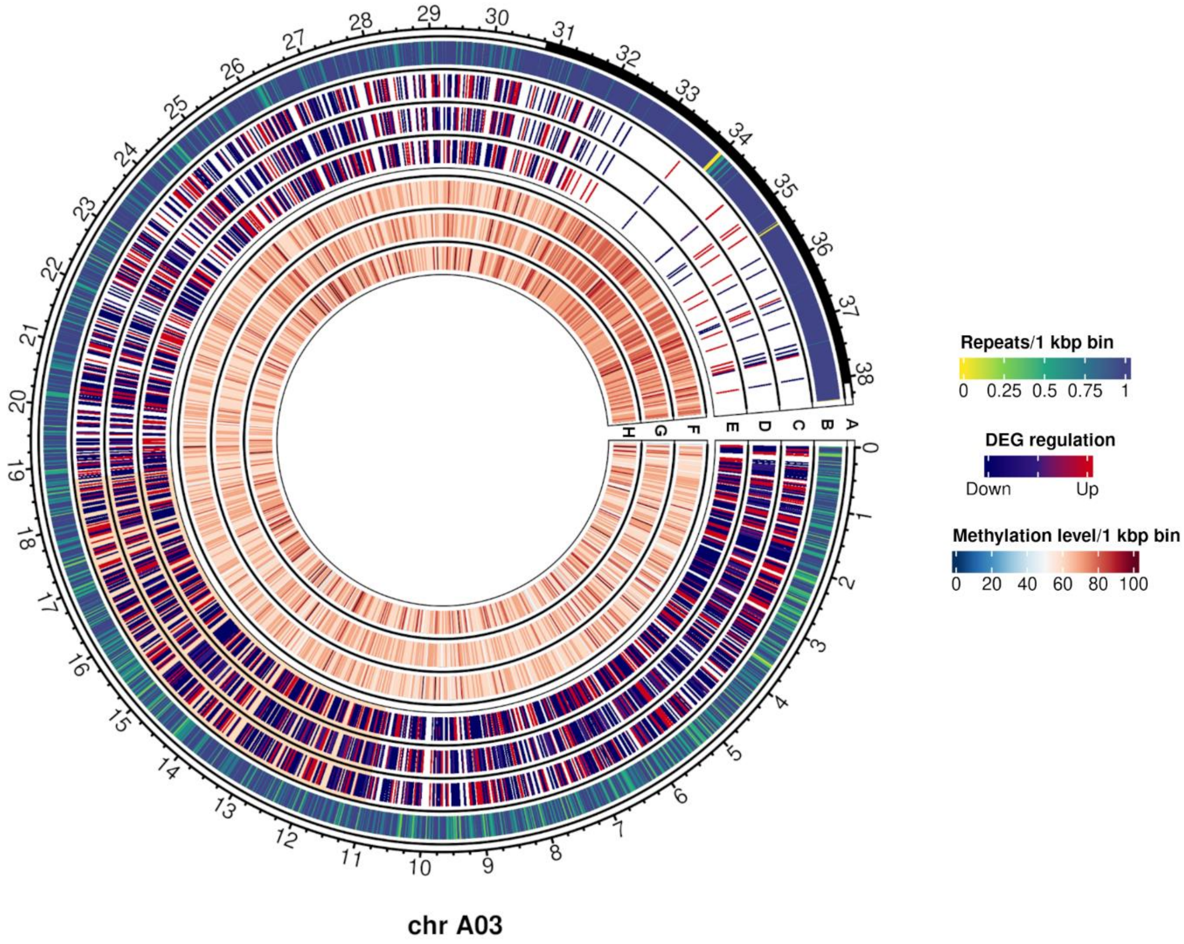


**Supplementary Figure 22** Differentially expressed genes (DEGs) and methylation levels from 30 days after pollination ovules from F0 and parents in chromosome A03. Outer to inner tracks correspond to: **a** Predicted centromere positions in black; **b** Repeat density per 1 kbp bin; **c-e**, DEG regulation in **(c)** Express 617, **(d)** F0 and **(e)** G3D001; **f-h:** Methylation levels per 1 kbp bin in **(f)** Express 617, **(g)** F0 and **(h)** G3D001. A differentially expressed chromosome segment between around 11 Mbp and 18.8 Mbp is highlighted in orange in tracks c-e.

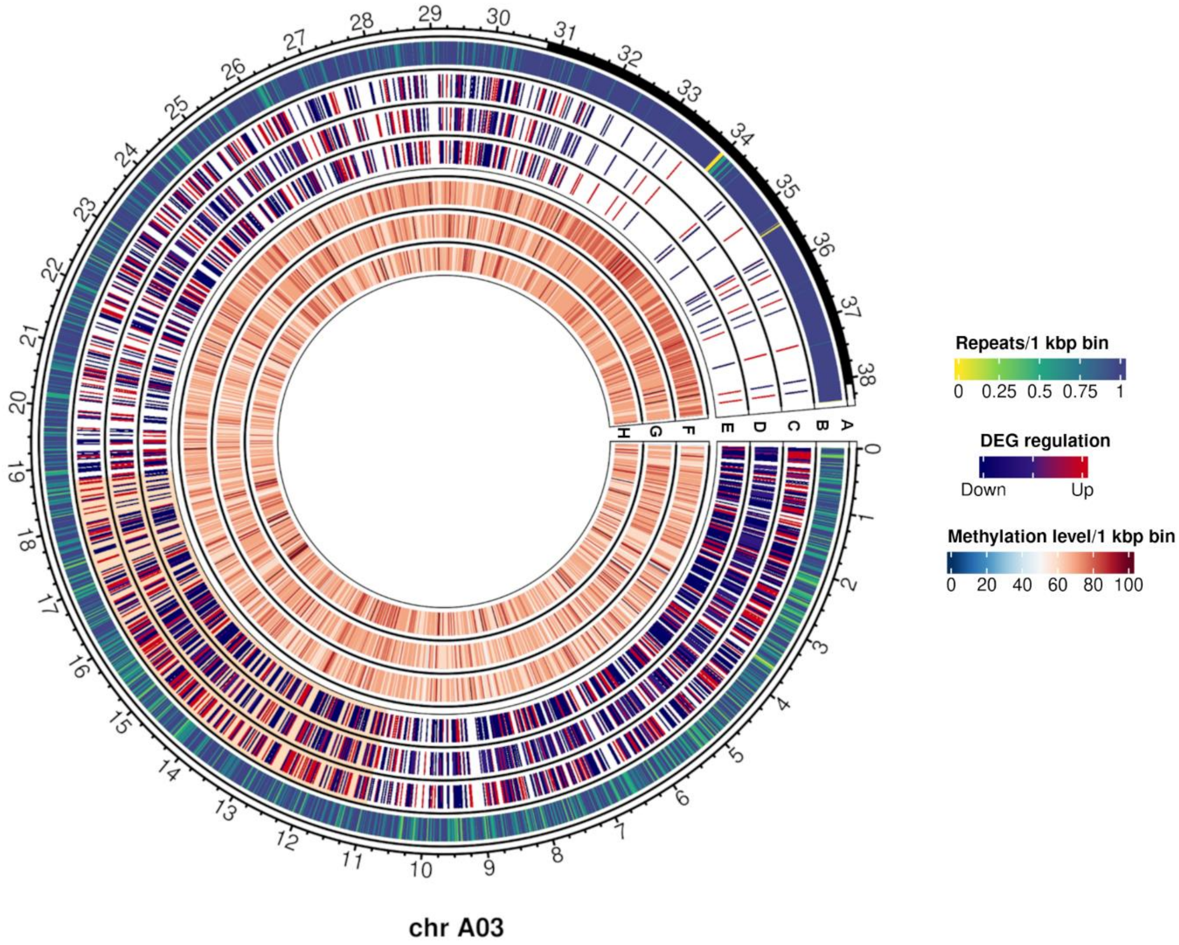


**Supplementary Figure 23** Differentially expressed genes (DEGs) and methylation levels from 15 days after pollination ovules from F1 and parents in chromosome A03. Outer to inner tracks correspond to: **a** Predicted centromere positions in black; **b** Repeat density per 1 kbp bin; **c-e**, DEG regulation in **(c)** Express 617, **(d)** F1 and **(e)** G3D001; **f-h:** Methylation levels per 1 kbp bin in **(f)** Express 617, **(g)** F1 and **(h)** G3D001. A differentially expressed chromosome segment between around 11 Mbp and 18.8 Mbp is highlighted in orange in tracks c-e.

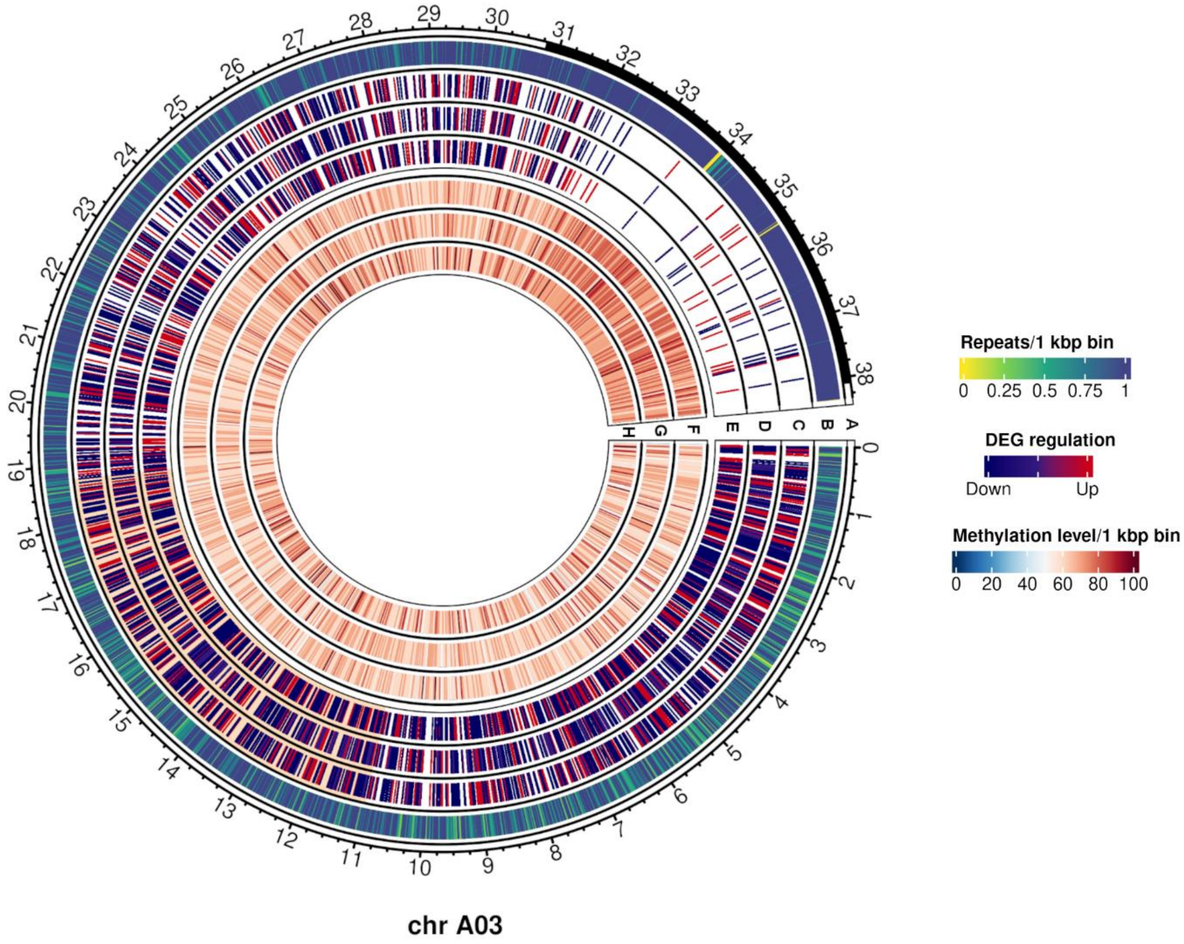


**Supplementary Figure 24** Differentially expressed genes (DEGs) and methylation levels from 30 days after pollination ovules from F1 and parents in chromosome A03. Outer to inner tracks correspond to: **a** Predicted centromere positions in black; **b** Repeat density per 1 kbp bin; **c-e**, DEG regulation in **(c)** Express 617, **(d)** F1 and **(e)** G3D001; **f-h:** Methylation levels per 1 kbp bin in **(f)** Express 617, **(g)** F1 and **(h)** G3D001. A differentially expressed chromosome segment between around 11 Mbp and 18.8 Mbp is highlighted in orange in tracks c-e.
